# Models for the architecture of the human inner kinetochore CCAN complex on centromeric α-satellite CENP-A nucleosome arrays

**DOI:** 10.1101/2025.11.04.686567

**Authors:** Cong Yu, Kyle W. Muir, Jing Yang, Ziguo Zhang, Stephen H. McLaughlin, David Barford

## Abstract

Human kinetochores assemble onto centromeric DNA defined by tandem copies of thousands of the 171 bp α-satellite repeat sequence. The centromere-specific CENP-A nucleosome (CENP-A^Nuc^) recruits the inner kinetochore constitutive centromere associated network (CCAN) complex. A previous cryo-EM structure of CCAN bound to a CENP-A^Nuc^ reconstituted with a 171 bp α-satellite repeat sequence showed how extranucleosomal linker DNA threads through an internal tunnel in the CCAN complex. To understand the higher-order architecture of the inner kinetochore assembled onto α-satellite repeat arrays, we have determined cryo-EM structures of CCAN with longer DNA sequences. These include free DNA and single and dimeric α-satellite repeats with CENP-A nucleosomes. We show from the structures of CCAN bound to both free DNA and monomeric CENP-A^Nuc^ that CCAN engages 65-70 bp of DNA comprising 30-35 bp of an upstream α-satellite repeat. This upstream DNA interacts with the histone-fold domain subunits of the CENP-TWSX module in a manner resembling how DNA is wrapped in nucleosomes. A complex of CCAN assembled onto a dimeric α-satellite repeat with two CENP-A nucleosomes showed that CCAN can only be accommodated on the linker DNA by unwrapping DNA from the CENP-TWSX module together with 25 bp of DNA from the upstream nucleosome. We discuss the implications of these results for models of CCAN assembly on arrays of α-satellite chromatin containing CENP-A^Nuc^.

## Introduction

The faithful inheritance of genetic information is crucial for life. Central to this process in eukaryotes are the roles fulfilled by kinetochores – large protein complexes that assemble at centromeric chromatin. Kinetochores function to attach sister chromatids to the mitotic spindle, and couple the work of microtubule depolymerization to chromatid segregation ^1–3^. Kinetochores also act as signalling hubs for the error correction and spindle assembly checkpoints that together coordinate the correct attachment of sister chromatids to the mitotic spindle with the initiation of sister chromatid segregation ^4,5^.

Human regional centromeres are megabase-sized structures, comprising thousands of contiguous 171 bp α-satellite repeat sequences. Almost identical α-satellite repeats are organized into higher order repeat (HOR) arrays of typically four, seven and eleven repeats ^6^. Only a small fraction of α-satellite repeats is associated with the approximately two-hundred CENP-A nucleosomes per centromere ^7^. CENP-A nucleosomes are restricted to the hypo-methylated centromere dip region (CDR), the most recently evolved segment of centromeres ^6,8,9^, and are interspersed with canonical H3 nucleosomes ^10^. Assuming at most 147 bp of DNA wraps a histone octamer ^11^, at least 24 bp of α-satellite repeat sequence forms extranucleosomal linker DNA. Both *in vivo* ChIP-seq mapping and in *vitro* reconstitution of CENP-A nucleosomes on α-satellite sequences indicate that the CENP-A nucleosome is installed precisely towards the 3’ end of the repeat, with the centre of the α-satellite repeat palindrome coinciding with the nucleosome dyad axis ^12,13^. Human centromeres, except for the Y chromosome centromere, contain the 17 bp B-box element. The majority of B-boxes are arranged on alternating α-satellite repeats ^14^ positioned such that the B-box on linker DNA partially overlaps a nucleosome entry site ^15^. CENP-B engagement by the B-box facilitates unwrapping of the DNA gyre from the CENP-A histone octamer ^16^, whereas the engagement of the dimeric CENP-B to two separate B-boxes induces looping of centromeric DNA ^17^.

The centromere-kinetochore assembly of vertebrates comprises large macromolecular complexes at the constriction point of chromosomes visible by electron microscopy in prometaphase cells as plate-like structures ∼200 nm in diameter, and attached to 20-30 microtubule filaments ^18^. Kinetochores are composed of two main components: the inner kinetochore centromere-binding CCAN complex, and the microtubule-binding KMN complex of the outer kinetochore ^19^. The 16-subunit CCAN complex is organized into five modules: CENP-C and CENP-OPQUR, and the DNA-binding modules, CENP-LN, CENP-HIKM and CENP-TWSX. The latter module comprises four histone-fold domain proteins ^20^. Cryo-EM structures of human and budding yeast CCAN have elucidated mechanisms of centromere and CENP-A nucleosome recognition ^12,21–25^. CCAN encloses extranucleosomal linker DNA within a topologically-enclosed chamber ^12,22^, a mechanism recently shown to be conserved in the CENP-A-independent holocentric kinetochores of *Lepidoptera* ^26^. In vertebrates, interactions with the CENP-A nucleosome, that confer CCAN specificity for CENP-A rather than H3 nucleosomes, are mediated primarily by the CENP-C subunit through its short linear interaction motifs that bridge the CENP-A histones of CENP-A^Nuc^ with the CCAN modules CENP-LN and CENP-HIKM ^12,21,23,27–30^. Human CENP-C incorporates two CENP-A-binding motifs, generating four potential CENP-A-binding sites (two CENP-A nucleosomes) in the CENP-C dimer ^31^.

In our previous cryo-EM structures of CCAN:CENP-A^Nuc^ and CCAN:DNA complexes, a DNA duplex was observed threading through a 50 Å-long central tunnel running the length of the CCAN molecule ^12^. This DNA-binding tunnel is formed by the CENP-LN channel, and CENP-I and CENP-TW subunits. As the DNA exits the tunnel it curves around the CENP-TWSX module, interacting with the histone-fold domains of CENP-TW. The DNA molecules used in the earlier study (the 171 bp α-satellite DNA used to reconstitute the CENP-A^Nuc^ for the CCAN:CENP-A^Nuc^ complex, and the 54 bp DNA used in the CCAN:DNA complex) were of insufficient length to extend beyond CENP-TW, and so in these complexes DNA did not contact CENP-SX of the histone-fold CENP-TWSX module. Previous cell biology and biochemical studies however, had identified CENP-SX as DNA-binding proteins at the centromere ^32^. We were therefore missing important details regarding how CCAN interacts with DNA at the centromere, and this information is required for understanding the higher-order architecture of kinetochores arranged on α-satellite repeat arrays. Furthermore, the repeating CCAN:CENP-A nucleosome unit at centromeres is not defined. Thus, the aims of this study are two-fold. First, we aim to obtain a more complete understanding of how CCAN interacts with DNA, and second we seek to provide models for how multiple copies of CCAN and CENP-A nucleosomes assemble onto contiguous α-satellite repeat sequences at human centromeres.

## Results

### CCAN binds 70 bp of DNA in a loop configuration that wraps the CENP-TWSX module

To understand how centromeric DNA interacts with CCAN, including all of the histone-fold domains (HFDs) of the CENP-TWSX module, we reconstituted CCAN with a longer DNA duplex (native human α-satellite DNA: 171 bp) and used mild cross-linking to stabilize the complex. The cryo-EM structure, resolved to 3.5 Å resolution, showed clearly defined cryo-EM density for ∼70 bp of DNA that wraps around CCAN in a loop-like structure (**Fig. 1 and Supplementary Figs 1 and 2a and Supplementary Table 1**). As observed in our previous CCAN:CENP-A^Nuc^ and CCAN:DNA complexes ^12^, DNA threads through the enclosed CCAN chamber, contacting CENP-TW as it exits the tunnel (**Fig. 1 and Supplementary Fig. 2a, b**). However, in the new structure, we observed an additional 20 bp of DNA exiting the DNA-binding tunnel, so that in total 40 bp of DNA interacts with the entire CENP-TWSX module. Additionally, a short segment of DNA (∼5 bp) extends beyond the CENP-TWSX module and approaches CENP-QU.

**Fig. 1.**
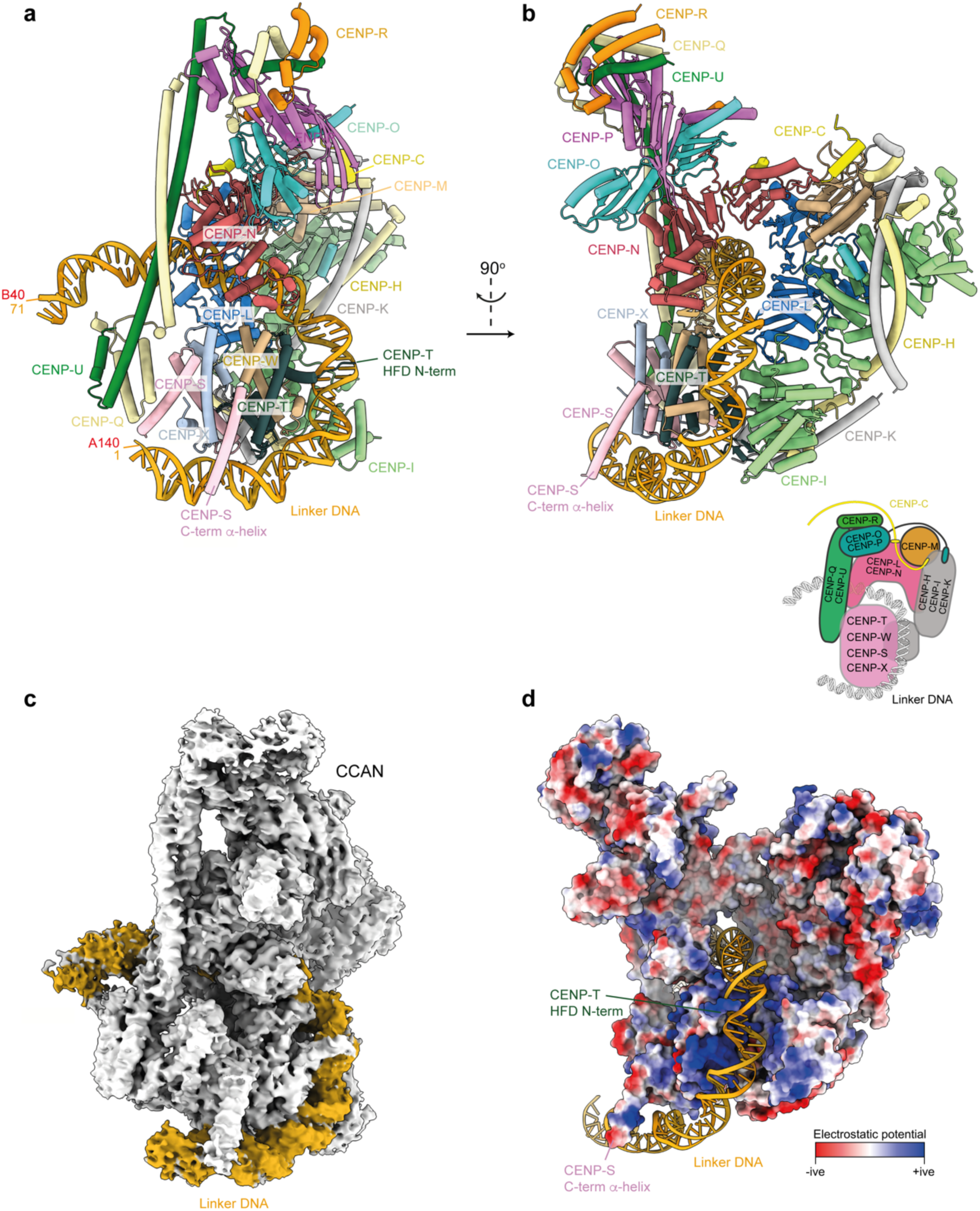
CCAN binds 70 bp of DNA in a loop configuration. **a** and **b**, Orthogonal views of the CCAN:DNA complex. 41 bp of DNA interacts with the CENP-TWSX module. Insert: Schematic of the CCAN:DNA complex. In panel (**a**), DNA is numbered ‘1’ and ‘71’ at the ordered 5’ and 3’ ends, and ‘A140’ and ‘B40’ to indicate the position of the DNA on a dimeric α-satellite sequence (Fig. 3c). **c**, Cryo-EM map of the CCAN:DNA complex with DNA coloured in gold and CCAN in grey. An unsharpened version of this map is shown in Supplementary Fig. 1c. **d**, Electrostatic surface of CCAN as viewed in panel B, with DNA shown as a ribbons representation (Supplementary Fig. 2).

**Fig. 2.**
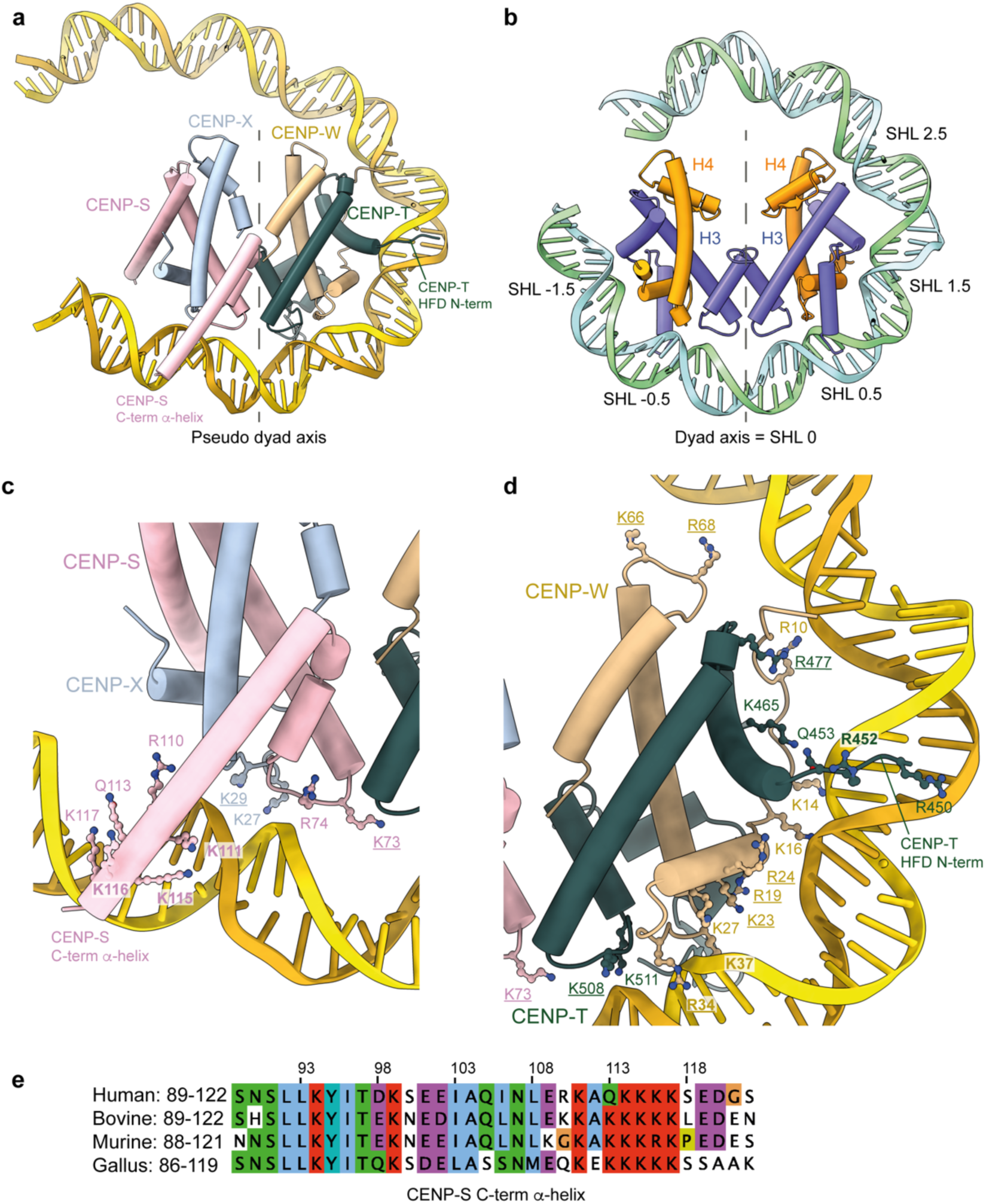
The CENP-TWSX module binds DNA analogous to the H3:H4 histone tetramer of canonical nucleosomes. **a**, CENP-TWSX and CCAN-associated 70 bp DNA, 40 bp interacts with the CENP-TWSX module. **b**, H3:H4 tetramer of a nucleosome with almost with 67 bp of DNA. Super helical locations (SHLs) are indicated (PDB:1AOI ^11^). The pseudo dyad axes of CENP-TWSX and the H3:H4 tetramer are shown in panels (a) and (b), respectively. **c**, Zoomed-view of panel (a) showing details of basic residues from the HFDs of CENP-S and CENP-X and CENP-S C-terminal α-helix interacting with the DNA phosphate backbone. **d**, Zoomed-view of panel (a) showing details of basic residues from the HFDs of CENP-T and CENP-W, and CENP-T N-terminal extension interacting with the DNA phosphate backbone. In panels (c) and (d) residues previously shown to contribute to CENP-TWSX – DNA binding are underlined ^32^. **e**, Multiple sequence alignment of the CENP-S C-terminal α-helix. Alignment created using Jalview ^33^.

The HFDs of the CENP-TWSX module assemble into a heterotetramer that structurally resembles the H3:H4 tetramer of canonical nucleosomes (**Fig. 2a, b**) ^11,32^. In the CCAN:DNA complex, DNA wraps around CENP-TWSX comparably to how H3:H4 tetramers wrap DNA in the context of an octameric nucleosome ^11^. In both structures DNA has a similar curvature and adopts a left-handed helical geometry. The CENP-TWSX module creates a positively-charged ridge that binds the DNA phosphate backbone (**Fig. 1d**). Strikingly, DNA wraps the H3:H4 and CENP-TWSX tetramers with identical registers such that the DNA major groove projects towards both the dyad axis of the H3:H4 tetramer, and the related pseudo-dyad axis of CENP-TWSX (**Fig. 2a, b**). Consequently, in both structures, at superhelical locations (SHLs) ±0.5, ±1.5, ±2.5, the DNA minor groove projects towards and contacts equivalent residues in the histone-fold domains of the H3:H4 tetramer and CENP-TWSX module. Our structure shows that DNA interacts with conserved basic residues of all four HFD domains of CENP-TWSX (**Fig. 2c, d**), consistent with prior mutagenesis data demonstrating the role of basic CENP-TWSX amino acids in DNA binding *in vitro* and centromere localization *in vivo* ^32^.

Additional protein-DNA contacts stabilize CENP-TWSX – DNA interactions. These involve an extended basic segment immediately N-terminal of the CENP-T HFD that inserts into the DNA minor groove, and a C-terminus α-helix of CENP-S that engages the DNA major groove (**Fig. 2a, c and d**). This CENP-S α-helix becomes ordered on binding DNA, such that a conserved stretch of basic residues stabilize its interactions with the DNA backbone (**Fig. 2c, e and Supplementary Fig. 2c, d**), as similarly observed in the crystal structure of a CENP-SX:DNA complex ^34^. Our structural data are consistent with previous studies showing that deletion of the C-terminus of CENP-S, including this basic region, abolished CENP-S binding to DNA *in vitro* ^32^. Similar observations regarding the interaction of linker DNA with the CENP-TWSX module were obtained by Musacchio and colleagues from a cryo-EM structure of a CCAN:DNA complex (M. Pesenti, I. Vetter and A. Musacchio, unpublished data).

### DNA of CENP-A^Nuc^ reconstituted with longer α-satellite repeat DNA wraps CENP-TWSX

We next assessed whether CENP-TWSX wraps linker DNA in the context of a CENP-A nucleosome. We assembled CCAN onto a CENP-A^Nuc^ reconstituted with 211 bp DNA that comprised the 171 bp α-satellite repeat and 40 bp from the 5’ upstream α-satellite repeat, and determined its structure using cryo-EM to an overall resolution of 4.5 Å (**Supplementary Fig. 3**). Both individual CCAN:DNA and CENP-A^Nuc^ modules were resolved in cryo-EM density, showing relative flexibility (**Supplementary Fig 3d**). A total of 200 bp DNA was visible in the cryo-EM map. Similar to our previous CCAN:CENP-A^Nuc^ structure with 171 bp DNA ^12^, CENP-A^Nuc^ is positioned at the 3’ end of the α-satellite repeat (**Fig. 3a, c and Supplementary Fig. 4a**). The 5’ terminus of the CENP-A^Nuc^ DNA gyre is unwrapped by 10 bp (**Fig. 3a and Supplementary Fig. 4b**). This unwrapped DNA then leads into extranucleosomal linker DNA that threads through the CCAN DNA-binding tunnel. On exiting the tunnel, DNA wraps around the entire CENP-TWSX module, as seen in the CCAN:DNA complex described above. This DNA comprises linker DNA of the 3’ downstream α-satellite repeat together with 30 bp of the 5’ upstream α-satellite repeat. Thus, a total of ∼65 bp of free DNA 5’ of CENP-A^Nuc^ interacts with CCAN (**Fig. 3c**). An unexpected finding was that compared with our previous CCAN:CENP-A^Nuc^ cryo-EM structure ^12^, CENP-A^Nuc^ adopts a different orientation relative to CCAN. In our new structure, CENP-A^Nuc^ is rotated by 216° and shifted by six bp away from CCAN. A shift of register while maintaining the same DNA phosphate backbone interactions with the CCAN DNA-binding tunnel causes the DNA duplex (and hence CENP-A^Nuc^) to rotate 36° per bp, rotating 216° overall (**Fig. 3b and Supplementary Video 1**). The reasons for the differences in CENP-A^Nuc^ position between the two CCAN:CENP-A^Nuc^ complexes are not entirely clear. A possible explanation is that the longer extranucleosomal DNA linker used in this study optimizes interactions with DNA-binding sites on CCAN, resulting in displacement of CENP-A^Nuc^. Cryo-EM density for CENP-C is observed at the CENP-C-binding sites on CENP-A^Nuc^, CENP-LN and CENP-HIKM, however, due to the displaced CENP-A^Nuc^, there are no direct contacts between CENP-A^Nuc^ and the back-side of CCAN.

**Fig. 3.**
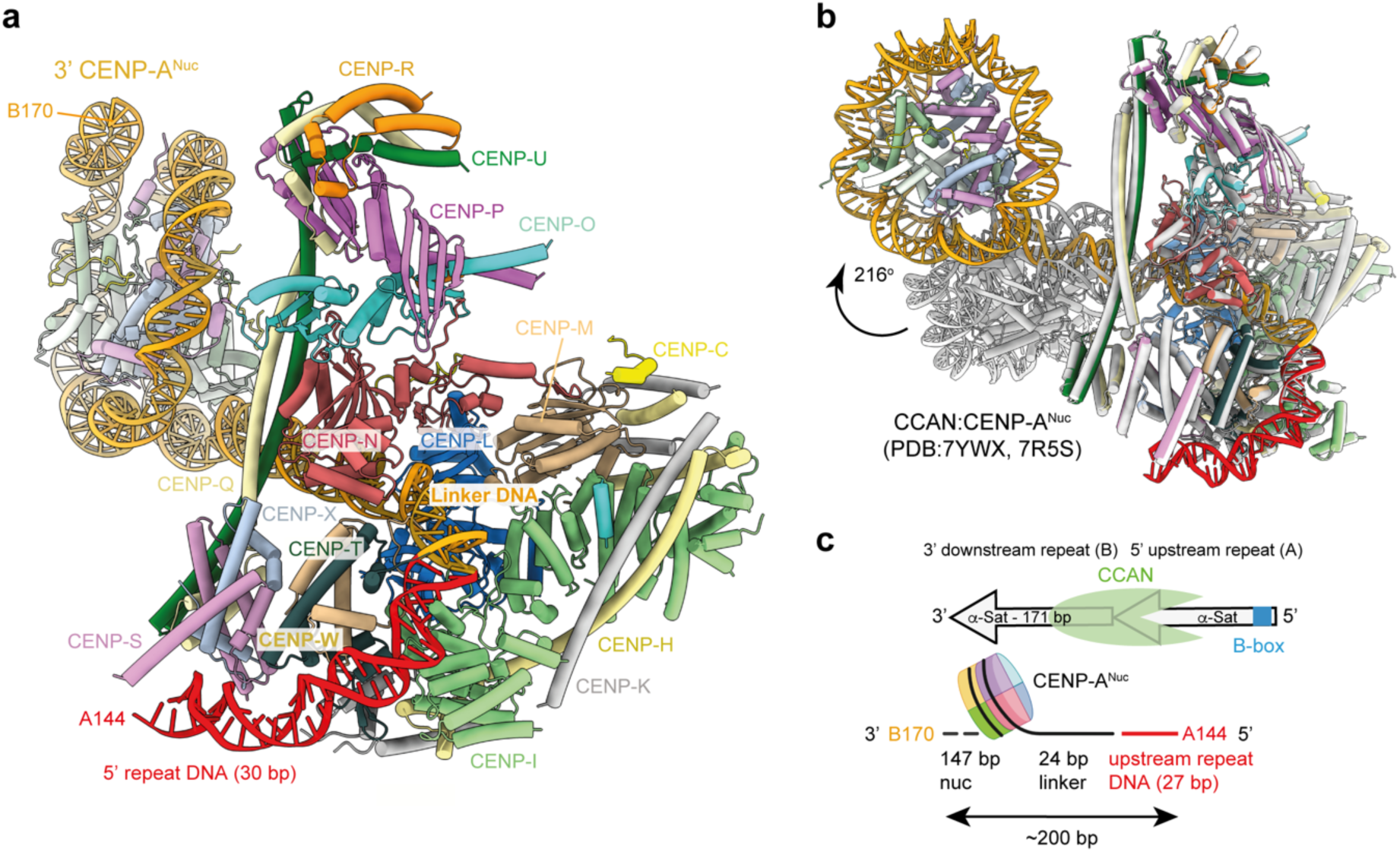
Cryo-EM structure of CCAN:CENP-A^Nuc^ showing CENP-TWSX binding DNA. **a**, Ribbons representation of the CCAN:CENP-A^Nuc^ complex. DNA coloured in red indicates DNA of the 5’ upstream α-satellite repeat. **b**, Superimposition of our new CCAN:CENP-A^Nuc^ complex (with 211 bp DNA) (coloured as in panel A) onto the previous model of CCAN:CENP-A^Nuc^ with a single 171 bp α-satellite repeat DNA (coloured grey) ^12^. This shows the difference in position and orientation of the downstream 3’ CENP-A^Nuc^. The two nucleosomes differ in orientation by 216° due to a 6 bp translation of the repeat sequence through the tunnel (**Supplementary Video 1**). **c**, Schematic of the CCAN:CENP-A^Nuc^ on an α-satellite repeat dimer. A total of ∼200 bp of DNA is embedded by the complex including 27 bp from the 5’ upstream α-satellite repeat. A B-box accessible for CENP-B binding in the upstream repeat is shown in blue. The nucleotide numbering (A144 and B170 in panels a and c) refers to the position of the visible DNA 5’ and 3’ ends of the complex within the α-satellite sequence. Positions in the α-satellite repeat are numbered 1-171 (5’ to 3’), and the upstream and downstream α-satellite repeats are labelled A and B, respectively (Supplementary Fig. 4).

Although the cryo-EM density for the DNA phosphate backbone is well resolved for both the CCAN:DNA and CCAN:CENP-A^Nuc^ complexes, density for individual DNA bases is less well defined such that purine and pyrimidine bases are not distinguished (**Supplementary Figs 1e, 2a and 3d**). We also do not observe sequence-specific protein-DNA interactions, which might be expected to contribute to a defined position of α-satellite DNA bound to CCAN. This poorly-resolved base-pair density is likely due to the limited resolutions of the CCAN:DNA and CCAN:CENP-A^Nuc^ complexes (3.5 and 4.5 Å, respectively), but would also be consistent with sequence variability at each position of the DNA duplex. However, the well-resolved cryo-EM density for the DNA phosphate backbone indicates that there is a specific mode of DNA recognition by CCAN, clearly illustrated by how the DNA duplex wraps around the CENP-TWSX module with phasing similar to how DNA wraps histone octamers (**Fig. 2a, b**).

In the CCAN:CENP-A^Nuc^ complex structure using the longer DNA described here, we do not observe contacts between CCAN and CENP-A^Nuc^, in contrast to our previous CCAN:CENP-A^Nuc^ complex ^12^. However, the position of CCAN on DNA appears fixed as judged by its defined position on the DNA duplex relative to CENP-A^Nuc^. This spacing of CCAN and CENP-A^Nuc^ is unlikely caused by the CENP-C subunit. First, the linker connecting the CENP-C binding sites on CENP-A^Nuc^ and CCAN (200 residues) is intrinsically disordered and not visible in the cryo-EM density. Second, CENP-C was also present in the previous structure with shorter DNA ^12^. Third, in the CCAN:di-CENP-A^Nuc^ complex without CENP-C (discussed below), the position of CCAN relative to the downstream nucleosome is the same as in this new CCAN:CENP-A^Nuc^ complex. In our previous structure ^12^, the 171 bp of a single α-satellite repeat DNA was too short to both wrap CENP-A^Nuc^ and extend beyond CENP-TW of the CENP-TWSX module. For that complex, to optimise CCAN-DNA interactions, CCAN would be required to shift towards CENP-A^Nuc^. In the new complex reported here, with 40 bp additional DNA (27 bp of which are ordered), there is sufficient DNA to wrap CENP-A^Nuc^, pass through the CCAN tunnel and wrap around CENP-TWSX. A fixed spacing between CENP-A^Nuc^ and CCAN positioned on DNA potentially indicates the presence of a CCAN-positioning sequence within α-satellite DNA that might contribute to how the DNA duplex wraps around the histone-fold CENP-TWSX module. This is reminiscent of how CENP-A nucleosomes are positioned precisely on α-satellite DNA, as revealed by *in vitro* reconstitution ^12^, and the phasing of CENP-A nucleosomes on centromeric chromatin ^13^, and of DNA nucleosome positioning sequences in general. The DNA-binding subunits of CCAN modules: CENP-LN, CENP-I, CENP-TWSX could all contribute to a defined mode of DNA phosphate backbone recognition.

### Implications for CCAN binding to α-satellite repeat dimers

Our new CCAN:DNA and CCAN:CENP-A^Nuc^ structures showing that CCAN itself interacts with 65 bp of duplex DNA has implications for understanding the higher-order organization of CCAN on repetitive α-satellite repeat CENP-A nucleosome arrays. The 65 bp DNA interacting with CCAN comprises 30 bp from the 5’ upstream α-satellite repeat and 35 bp of the 3’ downstream α-satellite repeat: 24 bp of extranucleosomal linker DNA and 10 bp of unwrapped CENP-A^Nuc^ DNA (**Fig. 3c**). The 30 bp DNA interacting with CCAN from the 5’ upstream α-satellite repeat is DNA of the CENP-A nucleosome gyre when CENP-A nucleosomes are positioned at the 3’ end of the α-satellite repeat ^12,13^. This suggests that in the context of a CCAN:CENP-A^Nuc^ complex, the upstream α-satellite repeat might be devoid of a nucleosome (either CENP-A or canonical H3). In alternative non-exclusive scenarios, to accommodate CCAN bound between two CENP-A nucleosomes on adjacent α-satellite repeats, a CENP-A nucleosome positioned on the upstream 5’ α-satellite repeat might unwrap by 30 bp, or the upstream CENP-A^Nuc^ may force DNA to unwrap from the CENP-TWSX module. A third possibility is that CCAN repositions CENP-A^Nuc^ by causing DNA to slide around the histone octamer. However, the defined position of CENP-A^Nuc^ at the 3’ of α-satellite repeats *in vivo* ^13^ does not support this scenario.

Substantial evidence indicates that the DNA termini of CENP-A nucleosomes are loosely wrapped and flexible. First, nuclease digestion sensitivity assays showed that CENP-A nucleosomes protect only 100-120 bp DNA, compared to 140-150 bp for canonical H3 nucleosomes ^13,24,35,36^. Second, FRET data indicated increased flexibility of terminal DNA segments in CENP-A^Nuc^ relative to canonical nucleosomes ^16^. Third, both crystal and cryo-EM structures of CENP-A^Nuc^ revealed unwrapping of terminal DNA segments ^12,16,24,37–41^, in some instances as much as 24-36 bp of flexible DNA at one end ^42^. Finally, CENP-B and CENP-C have been implicated in promoting DNA unwrapping from CENP-A^Nuc^ (Refs ^16,43,44^).

### CCAN binds a dimeric α-satellite repeat with two CENP-A nucleosomes

To investigate the feasibility of CCAN binding to linker DNA connecting two adjacent CENP-A nucleosomes, we reconstituted two CENP-A nucleosomes on an α-satellite repeat dimer. The α-satellite repeat dimer was designed to include only linker DNA connecting the two CENP-A nucleosomes: upstream and downstream α-satellite repeats were 147 bp and 171 bp, respectively (**Supplementary Fig. 5a**). To stabilize the two nucleosome positions on this 324 bp DNA, we replaced 92 bp of native α-satellite repeat DNA at the centre of each CENP-A^Nuc^ site with Widom601 DNA. We attempted to reconstitute CENP-A nucleosomes on a dimeric α-satellite repeat sequence (342 bp) using completely native sequence. Whereas we could generate well-positioned CENP-A nucleosomes on a single α-satellite repeat (as defined by native agarose gels), we found that native sequences exceeding 180-190 bp did not wrap well. Typically, the shifts were incomplete and heterogeneous with substantial amounts of free DNA compared to monomeric α-satellite DNA and Widom-based sequences.

We observed that CCAN associated with this di-CENP-A^Nuc^ array, as assessed by size exclusion chromatography (SEC) (**Supplementary Fig. 5b, c**). Analysis of the peak SEC fractions using interferometric scattering microscopy (iSCAT) revealed two species: the major 1,054 kDa complex corresponded to a single CCAN protomer bound to the di-CENP-A^Nuc^ array (CCAN:di-CENP-A^Nuc^), although with a lower mass than the expected 1,282 kDa. The underestimate of molecular mass using ISCAT may be due to the DNA component which scatters less than the globular proteins used for calibration. A 445 kDa component corresponded to a di-CENP-A^Nuc^ array without CCAN (expected molecular mass of 431 kDa) (**Supplementary Fig. 5d**). The partial dissociation of CCAN from di-CENP-A^Nuc^ is likely due to the low sample concentrations required for single molecule counting in iSCAT.

#### CCAN and CENP-B binding to the same linker DNA segment is mutually exclusive

Our α-satellite dimer incorporated a B-box within the linker DNA of the downstream α-satellite repeat connecting the two CENP-A nucleosomes (**Supplementary Fig. 6a**). SEC data showed that the CENP-B protein recognized the B-box specifically because CENP-B bound to the di-CENP-A^Nuc^ array with an intact B-box, but not to a di-CENP-A^Nuc^ array with a mutant B-box (**Supplementary Fig. 6b, c**). We next tested whether CCAN and CENP-B bind the di-CENP-A^Nuc^ array simultaneously. We observed that CCAN displaced CENP-B from the array, showing that CCAN and CENP-B engagement of linker DNA is mutually exclusive (**Supplementary Fig. 6d, e**). This is consistent with our model of CCAN:CENP-A^Nuc^, that with CCAN bound to linker DNA, the B-box is enclosed by the CCAN DNA-binding tunnel and therefore inaccessible to CENP-B.

### CCAN binds to linker DNA in the dimeric α-satellite repeat with two CENP-A nucleosomes

To understand the structural basis of how CCAN interacts with linker DNA connecting adjacent CENP-A nucleosomes, we prepared cryo-EM grids with the CCAN:di-CENP-A^Nuc^ complex. Despite cross-linking and the use of additives to reduce sample denaturation at the air-water interface, CCAN:di-CENP-A^Nuc^ particles were not visible on either cryo-electron micrographs or subsequent 2D classes of the cryo-EM dataset. We instead observed a heterogeneous mixture of free DNA, CENP-A^Nuc^ and protein complexes (data not shown), indicative of CCAN:di-CENP-A^Nuc^ denaturation on cryo-EM grids. To stabilize the CCAN:di-CENP-A^Nuc^ complex for cryo-EM analysis, we combined it with the CENP-A nucleosome stabilizing-single chain antibody fragment (ScFv) ^40^, omitting CENP-C due to its binding site overlapping that of the ScFv (**Supplementary Fig. 7)**, and collected a large cryo-EM dataset on this ScFv-stabilized CCAN:di-CENP-A^Nuc^ complex (**Supplementary Figs 8 and 9 and Supplementary Table 1**).

2D classification of the selected particles and subsequent 3D reconstruction and classification revealed a highly heterogenous mixture of molecular states (**Supplementary Figs 8b-d and 9**). The majority of 2D classes from nearly nine million picked particles comprised either free mono-CENP-A nucleosome or CCAN:DNA with no obvious associated CENP-A nucleosome. *Ab initio* reconstruction and 3D refinement resulted in high resolution maps of CCAN:DNA and a CENP-A mono-nucleosome with bound ScFv (2.9 Å and 2.7 Å, respectively; **Supplementary Fig. 9**). The CCAN reconstruction showed DNA enclosed within the DNA-binding tunnel. We re-extracted CCAN particles in RELION ^45^ using a larger box and performed further 3D classification (**Supplementary Fig. 9**). This procedure revealed 3D classes with diffuse, poorly-defined EM densities at both 3’ and 5’ entrances to the CCAN-DNA tunnel that likely represented flexible CENP-A nucleosomes. To determine whether the density at the 3’ entrance corresponded to a 3’ CENP-A nucleosome, we generated a wide mask covering this density and performed signal-subtraction of density outside the mask. Subsequent 3D reconstruction and classification of the mask-generated recentred particles recovered a CENP-A nucleosome, confirming that the diffuse density observed in 2D classes as CENP-A^Nuc^. Using the corresponding unmasked particle set, we then calculated a new 3D reconstruction at 9.5 Å resolution that revealed both CCAN and 3’ CENP-A^Nuc^. Consistent with this reconstruction, 2D classes calculated from this particle set showed density features for CCAN and 3’ CENP-A^Nuc^ (**Supplementary Fig. 8c**). A similar approach was used to analyse EM density at the 5’ entrance to the CCAN-DNA tunnel, resulting in a 3D reconstruction with CCAN and associated 5’ CENP-A^Nuc^ at 15 Å resolution (**Supplementary Figs 8d and 9**). In the 2D class galleries corresponding to both the 3’ CCAN:CENP-A^Nuc^ and 5’ CCAN:CENP-A^Nuc^ complexes, we observed diffuse densities at the opposite end of the CCAN DNA-tunnel (i.e. 5’ and 3’, respectively), for example **Supplementary Fig. 9a(i)**. These particles are consistent with CCAN bound to the linker DNA of the di-CENP-A^Nuc^ array. However, the small number of the particles precluded a 3D reconstruction. We therefore generated a composite cryo-EM map of a CCAN:di-CENP-A^Nuc^ complex from the individual 3’ CCAN:CENP-A^Nuc^ and 5’ CCAN:CENP-A^Nuc^ maps (**Supplementary Fig. 9a(i)** and **Fig. 4b**). One of its projected views matches an experimental 2D class of a CCAN:di-CENP-A^Nuc^, validating our composite approach (**Supplementary Fig. 9a(i)**).

**Fig. 4.**
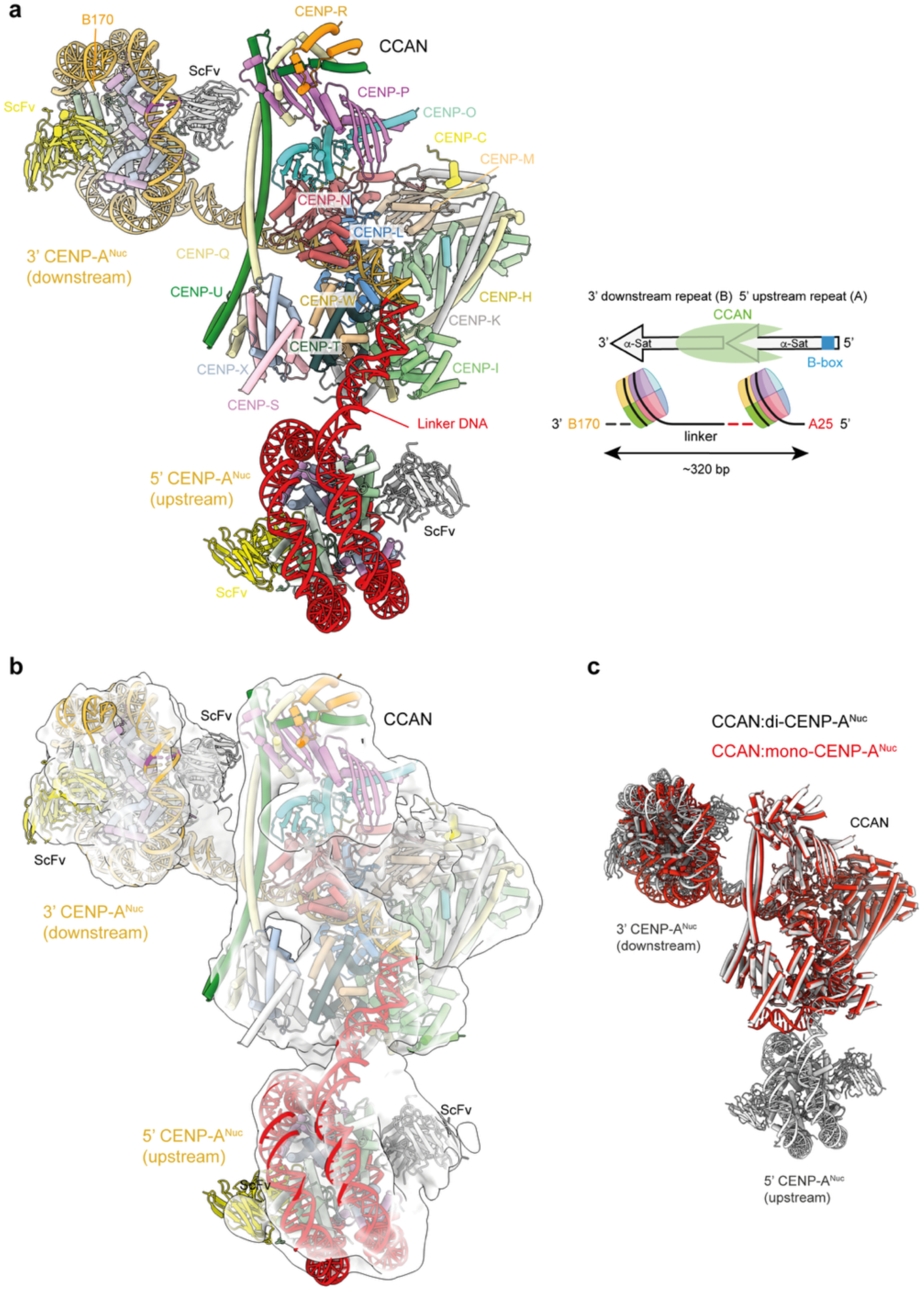
Structure of the CCAN:di-CENP-A^Nuc^ complex. **a**, Ribbons representation of CCAN:di-CENP-A^Nuc^ complex. Insert: Schematic of the CCAN:di-CENP-A^Nuc^ on a dimer of α-satellite repeats. 20-25 bp of the upstream 5’ CENP-A nucleosome is unwrapped. A total of 300-310 bp of DNA is embedded by the complex. A B-box accessible for CENP-B binding in the upstream repeat is shown in blue. **b**, Cryo-EM map of CCAN:di-CENP-A^Nuc^ complex. **C**. Superimposition of the new CCAN:CENP-A^Nuc^ and CCAN:di-CENP-A^Nuc^ complexes showing the identical position of the 3’ CENP-A^Nuc^. The nucleotide numbering (A25 and B170 in panel a) is defined in Fig. 3c (Supplementary Fig. 10).

#### CCAN bound to linker DNA forces DNA unwrapping from CENP-TWSX and 5’ CENP-A^Nuc^

5’ CCAN:CENP-A^Nuc^ complex: Despite the limited resolution of this map, reliable alignment of CENP-A^Nuc^ into cryo-EM density was facilitated by the ScFv associated with both sides of the nucleosome (**Fig. 4b**). Analysis of this structure indicated that a CENP-A nucleosome positioned 5’ to CCAN is incompatible with DNA wrapping around the CENP-TWSX module (**Fig. 4a, b and Supplementary Fig. 10a)**. Analogous to our previous CCAN:CENP-A^Nuc^ and CCAN:DNA complexes ^12^, we observed that as DNA exits the CCAN-DNA tunnel, it runs along the CENP-I-CENP-TW channel, although unlike our new CCAN:DNA and CCAN:CENP-A^Nuc^ complexes (this study), does not continue to bend around CENP-SX, but instead projects in a straighter trajectory to join the DNA of the 5’ CENP-A nucleosome. The DNA gyre of the 5’ CENP-A nucleosome connected to linker DNA is itself unwrapped by 20-25 bp (**Supplementary Fig. 10c**). Thus, even without DNA wrapping around CENP-SX, the 5’ CENP-A nucleosome is forced to unwrap to allow its positioning onto an upstream α-satellite repeat. Our fitting of the 5’ CENP-A^Nuc^ to the cryo-EM density map is consistent with the α-satellite repeat length of 171 bp such that the DNA length from the end of the downstream (3’) α-satellite repeat sequence to the dyad axis of the 5’ CENP-A^Nuc^ is 74 bp (**Fig. 4a**).

In the 3’ CCAN:CENP-A^Nuc^ complex, the position and orientation of the 3’ CENP-A^Nuc^ in the CCAN:di-CENP-A^Nuc^ complex is the same as our new CCAN:mono-CENP-A^Nuc^ complex with 211 bp DNA (this study), differing from that in the previously published CCAN:CENP-A^Nuc^ complex ^12^ (**Fig. 4a, c and Supplementary Fig. 10a, b**). Thus, the ScFv used for the CCAN:di-CENP-A^Nuc^ structure determination does not affect the orientation of the 3’ CENP-A^Nuc^, consistent with modelling showing that ScFv binding to CENP-A^Nuc^, when in the mono-CENP-A^Nuc^ position, is possible without a steric clash with CCAN.

## Discussion

In this study we investigated the molecular basis for how human CCAN interacts with CENP-A nucleosomes and DNA of α-satellite repeat arrays, revealing that CCAN engages as much as 70 bp of DNA. As described previously ^12^, DNA threads through the enclosed tunnel generated by the CENP-LN module, CENP-I and CENP-TW subunits. The longer DNA sequences used in this study showed additional DNA interactions with the CENP-SX subunits of the CENP-TWSX HFD module. The CENP-TWSX subunits interact with DNA analogously to how the H3:H4 heterotetramer, in the context of octameric nucleosomes, wraps its DNA gyre. The specific details of the interactions of DNA with CENP-SX, together with CENP-TW, observed in our structure, are in agreement with biochemical and cell biology data, confirming the proposal that human CCAN incorporates a nucleosome-like particle at the centromere ^20,32^. Consistent with this, CENP-TWSX augments the affinity of CCAN for a CENP-A nucleosome three-fold ^12^. Thus, this study indicates that CCAN creates a tight attachment to centromeric DNA by both topological entrapment of DNA through the CCAN DNA-binding tunnel (CENP-LN, CENP-I), and by wrapping DNA around a HFD tetramer (CENP-TWSX), mimicking the mechanism by which nucleosomes package DNA.

The finding that CCAN alone engages 65-70 bp of DNA has implications for the higher-order architecture of CCAN and CENP-A nucleosomes on repetitive α-satellite sequences. The 65 bp of extranucleosomal DNA bound by CCAN greatly exceeds the 24 bp linker DNA connecting two adjacent and fully wrapped nucleosomes (147 bp) on tandem α-satellite repeats. In our CCAN:CENP-A^Nuc^ structures, 10 bp of DNA are unwrapped from the 3’ CENP-A nucleosome gyre as it enters the CCAN DNA-binding tunnel (Ref. ^12^ and this study, **Supplementary Fig. 4b**). Thus, to accommodate 65 bp of DNA interacting with CCAN requires ∼30 bp DNA of the upstream α-satellite repeat, in addition to the 24 bp of α-satellite extranucleosomal linker DNA and 10 bp of unwrapped DNA from the downstream CENP-A nucleosome (**Fig. 3c**). Since CENP-A nucleosomes are positioned exactly on α-satellite repeats such that the 3’ end of the repeat defines the start of the CENP-A nucleosome DNA gyre ^12,13^, the 30 bp of upstream α-satellite repeat DNA would require the upstream CENP-A nucleosome to unwrap 30 bp of DNA. Such unwrapping is potentially feasible as demonstrated by the extensive levels of DNA unwrapping from CENP-A nucleosomes documented previously ^12,16,24,37–42^. To test whether CCAN associates with DNA connecting two adjacent CENP-A nucleosomes, we reconstituted a dimeric α-satellite repeat with CENP-A nucleosomes. SEC and iSCAT data confirmed interactions of CCAN with a dimeric α-satellite repeat with two CENP-A nucleosomes (**Supplementary Fig. 5**). That this interaction required the free linker DNA connecting the two nucleosomes was confirmed by the ability of CCAN to displace CENP-B from its B-box binding site located within the linker DNA (**Supplementary Fig. 6d, e**). Our cryo-EM structure of the CCAN:di-CENP-A^Nuc^ complex revealed CCAN bound to DNA linking two CENP-A nucleosomes (**Fig. 4**). However, accommodating CCAN at this site involved not only 20-25 bp of DNA to be unwrapped from the 5’ upstream CENP-A nucleosome, but also dissociation of DNA from CENP-SX. Thus, although the upstream CENP-A nucleosome is unwrapped, the available free DNA duplex is insufficient to bind CENP-SX, presumably because of steric hindrance between the upstream CENP-A nucleosome and CCAN.

Whether such an arrangement of tandem CENP-A nucleosomes and central CCAN that we observe in our CCAN:di-CENP-A^Nuc^ complex occurs at native human centromeres is unclear. The dissociation of DNA from CENP-SX is inconsistent with the role of conserved basic residues of CENP-SX in binding DNA revealed by our CCAN:DNA and CCAN:mono-CENP-A^Nuc^ structures and from prior *in vitro* binding assays, and the role of CENP-SX in mediating kinetochore assembly in cells ^32^. Furthermore, the extensive unwrapping of DNA from a CENP-A nucleosome, although observed for a small population of particles in a cryo-EM study ^42^, is not consistent with CENP-A nucleosomes defining a 120 bp footprint *in vivo* ^13,46^, and the size of the nuclease-resistant core observed *in vitro*. Although our cryo-EM structure of the CCAN:di-CENP-A^Nuc^ complex embeds a total of 300-310 bp (**Fig. 4a**), in agreement with the 250-300 bp centromere core-selective footprint identified by single molecule chromatin fibre sequencing (Fiber-seq) ^46^, we propose two alternative scenarios for the arrangement of CCAN and CENP-A nucleosomes on α-satellite arrays. In one scenario, the kinetochore unit would constitute a monomeric CCAN:CENP-A^Nuc^ complex in which DNA from the upstream α-satellite repeat engages CENP-TWSX such that a total of 200-210 bp of DNA is bound by the CCAN:CENP-A^Nuc^ complex (**Fig. 3**). In a second related scenario, a central CENP-A nucleosome would interact with two flanking CCAN protomers in a pseudo-symmetric arrangement of the mono-CCAN:CENP-A^Nuc^ complex (di-CCAN:CENP-A^Nuc^) (**Fig. 5**). This proposed assembly, with the central CENP-A nucleosome and associated CCAN protomers spanning three α-satellite repeats, embeds 260-270 bp. The 210 bp bound by mono-CCAN:CENP-A^Nuc^ is shorter than the 250-300 bp centromere core-selective footprint ^46^, however the 260-270 bp bound to the proposed di-CCAN:CENP-A^Nuc^ complexes matches the reported 250-300 bp centromere footprint. We note that Dubocanin et al., (2025) also identified centromere-associated footprints of 210 bp in size ^46^. Although these were assigned as di-nucleosomes, it is also plausible that these methylation-free footprints correspond to mono-CCAN:CENP-A^Nuc^ assemblies. Consistent with this, a recent cryo-ET study of human centromeres found that the majority of putative kinetochore particles were consistent in size with a mono-CCAN complex closely associated with a single CENP-A nucleosome ^47^. Additionally, larger globular densities containing multiple nucleosomes and CCAN complexes were also observed, although whether these constitute tandem CCAN and CENP-A^Nuc^ complexes was not defined.

**Fig. 5.**
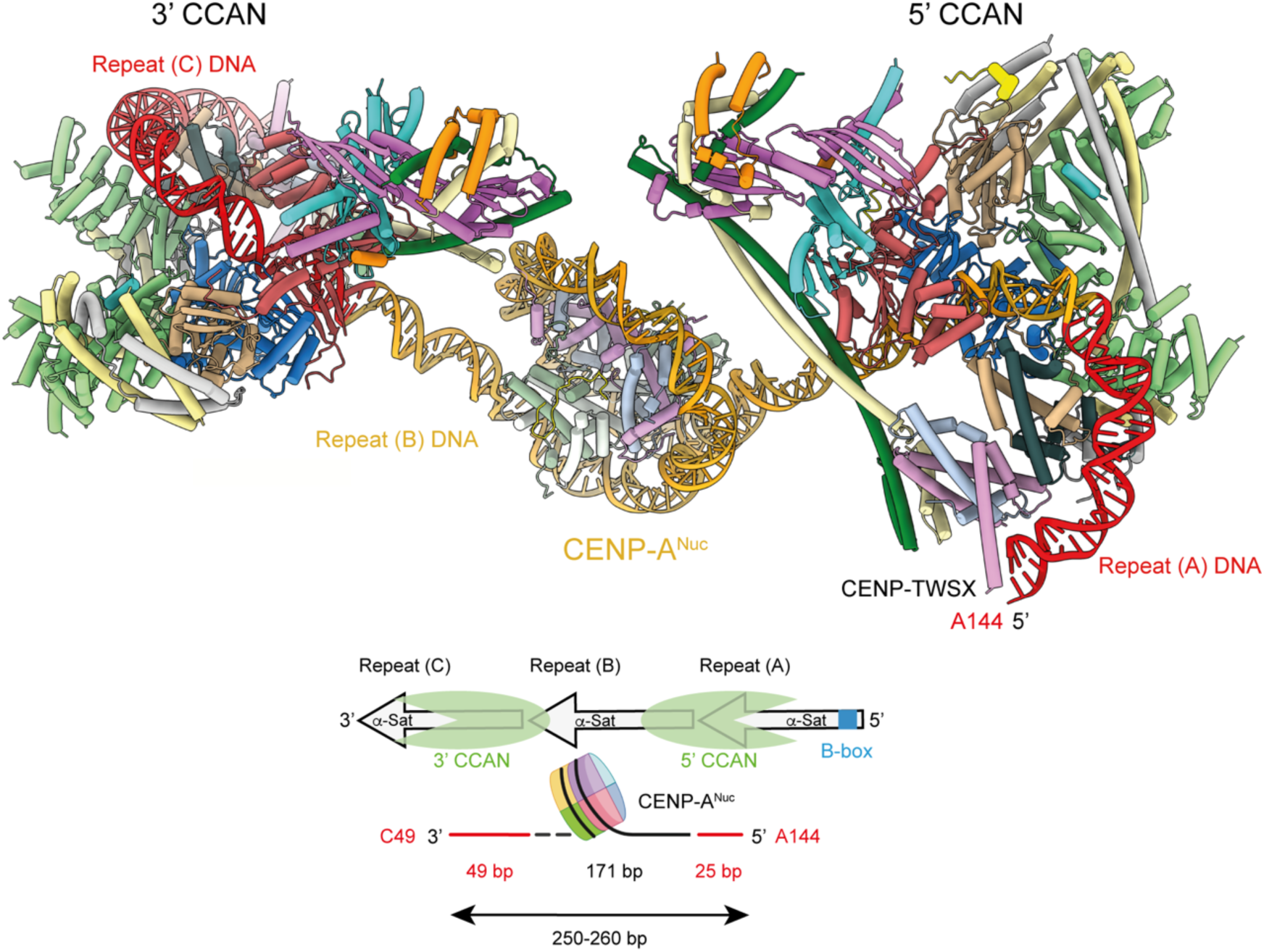
Model of a di-CCAN:CENP-A^Nuc^ complex. The model was generated from the CCAN:mono-CENP-A^Nuc^ structure by superimposing on CENP-A^Nuc^. The two CCANs together with a single CENP-A^Nuc^ overlap three α-satellite repeats. In the model, DNA from the central α-satellite sequence is coloured gold. DNA bound to the two CCANs from the 5’ and 3’ α-satellite repeats (repeats A and C, respectively) is coloured red. Insert: Schematic of the model with DNA coloured red for the 5’ and 3’ repeats as in the model. A B-box accessible for CENP-B binding in the upstream repeat is shown in blue. B-boxes in the two upstream repeats are occluded by CCAN. 260-270 bp of DNA are bound by the di-CCAN:CENP-A^Nuc^ complex. The nucleotide numbering (A144 and C49) is defined in Fig. 3.

Our models are consistent with the published numbers of CENP-A, CENP-C and CENP-T proteins at centromeres, and the region of the centromere occupied by CENP-A nucleosomes. Within canonical centromeres, the two-hundred CENP-A nucleosomes ^7^ are restricted to a region of ∼100-500 kilobases ^6,13^ that defines kinetochore assembly. Since 100-500 kilobase comprises 600-3,000 α-satellite repeats, this indicates one CENP-A nucleosome per 3-15 α-satellite repeats. The question of how many CENP-A nucleosomes a CENP-C dimer binds at the centromere is unclear. Since each human CENP-C subunit contains two-CENP-A binding motifs, in principle a CENP-C dimer could bind two CENP-A nucleosomes and two CCAN protomers, consistent with *in vitro* data ^31^. However, for human CENP-C, only one CENP-A binding motif is essential ^13,48–50^, and chicken, *Xenopus* and *S. cerevisiae* CENP-C all contain one CENP-A binding motif. For the human kinetochore, possible models are for one CENP-C dimer for either one or two CENP-A nucleosomes, and one or two CCAN protomers. Suzuki et al reported 215 copies of CENP-C (108 CENP-C dimers) and 72 copies of CENP-T ^51^ per centromere, data roughly consistent with the number of CENP-A nucleosomes. The three models of CCAN:CENP-A^Nuc^ assembly discussed here: (i) 1xCCAN:1xCENP-A^Nuc^:1xCENP-C dimer (**Fig. 3**), (ii) 1xCCAN:2xCENP-A^Nuc^:1xCENP-C dimer (**Fig. 4**), and (iii) 2xCCAN:1xCENP-A^Nuc^:1xCENP-C dimer (**Fig. 5**) are all feasible candidates given the published copies of CENP-A, CENP-C and CENP-T proteins per centromere. Models (i) and (ii) would occupy two α-satellite repeats, whereas model (iii) would occupy three repeats. Assuming 200 CENP-A nucleosomes per centromere and contiguous tandem repeats of the CCAN:CENP-A^Nuc^ assembly, the number of α-satellite repeats required is 400 (models (i) and (ii) or 800 (model (iii)), consistent with the 600-3,000 α-satellite repeats of the ∼100-500 kilobase CENP-A region of the centromere ^6,13^. Our models are thus consistent with the published numbers of CENP-A, CENP-C and CENP-T proteins, and the region of the centromere occupied by CENP-A nucleosomes.

In a model of a CENP-C dimer binding to a CCAN:di-CENP-A^Nuc^ complex, a possible scenario is for the pair of C-terminal CENP-A binding sites (CENP-C motif) to interact with either the upstream or down-stream CENP-A nucleosome and the pair of N-terminal CENP-A binding sites (central region) to interact with the remaining CENP-A nucleosome. As there is one CCAN protomer in this complex, only one CENP-C subunit of the dimer would bind CCAN. Chicken CENP-C dimers are observed to form higher-order oligomers based on the crystal packing of chicken CENP-C cupin dimers, supported by biochemical and cell biology data ^52^. The same study also presented evidence for oligomerization of human CENP-C cupin domains ^52^, as had been reported previously ^53^, although *in vitro* oligomers of cupin dimers were not observed ^31^. Formation of higher-order oligomers of CENP-C might be expected to oligomerize the CCAN:CENP-A^Nuc^ assemblies. However, we are unable to reliably predict the exact architecture of this higher-order assembly. Hara et al., (2023) ^52^ proposed that CENP-C oligomerization would lead to increased CENP-C concentration that might stabilize CCAN assembly.

Our findings that CCAN displaces CENP-B from B-box linker DNA indicated that the binding of CCAN and CENP-B to a single α-satellite repeat-CENP-A^Nuc^ module is mutually exclusive (**Supplementary Fig. 6**). B-boxes are generally present on alternating α-satellite repeats ^6^. For all three models of CCAN:CENP-A^Nuc^ complexes discussed above (**Figs 3-5**), a B-box in the upstream α-satellite repeat is accessible for CENP-B binding, although only in models (i) and (ii) would CENP-B bind to B-box elements on alternat α-satellite repeats.

In summary this work, combined with recent 2D and 3D mapping of CCAN, CENP-A nucleosomes and DNA-binding proteins at native centromeres ^46,47^, suggest potentially heterogenous structures of CCAN and CENP-A nucleosomes assembled on α-satellite repeat arrays including mono-CCAN:CENP-A^Nuc^, di-CCAN:CENP-A^Nuc^ and CCAN:di-CENP-A^Nuc^ complexes.

**Supplementary Video 1.** Video illustrates the difference in position of the 3’ CENP-A nucleosome between our previous CCAN:CENP-A^Nuc^ complex with 171 bp DNA (grey cartoon) ^12^ and the new structure of CCAN:CENP-A^Nuc^ with 211 bp DNA (ASW6) (colour-coded according to subunits as in Fig. 1). Relative to the previous CCAN:CENP-A^Nuc^ structure, in the new model, the DNA register of the α-satellite repeat is shifted 6 bp (translation of 20 Å) away from CCAN (towards the 3’ end). Because the DNA phosphate backbone contacts with CCAN are preserved, this results in a 216° rotation of CENP-A^Nuc^.

## Methods

### DNA cloning

Protein complexes for CCAN and CENP-A histone octamer were prepared as previously described ^12^, except CENP-Q was modified to CENP-Q^55–268^, CENP-U was modified to CENP-U^235-418^, CENP-I was changed to isoform 1 from previously isoform 2, and CENP-B full length was cloned into pOPINS plasmid, with a double strep II-sumo-TEV tag at their N-termini. The DNA fragment ASW14 was designed as two repeats from α-satellite DNA of chromosome 2 with 91 base pairs centred on their dyad positions replaced with Widom601 sequence aligned at the dyad base. A B-box sequence is in the linker region. Six ASW14 fragments flanked by GATATC (EcoRV recognition site) were assembled into the pMK-RQ plasmid. The plasmid containing ASW14 fragments was produced by using Plasmid Plus Giga Kit (QIAGEN). The DNA fragment was released by EcoRV-HF (NEB) at 37 °C for 16 h. The digested sample was diluted five-fold with PBS and loaded onto a pre-equilibrated (PBS) 1 mL Resource Q Column (GE 17-1177-01). The column was washed with 10 mL PBS (buffer A), followed by a wash with 20 mL of 50% of buffer B (PBS with 850 mM NaCl). The DNA fragment was eluted using a gradient to 70% of buffer B at 1 mL/min over 120 min. The target fragment eluted around 60% buffer B. The fractions containing DNA were checked on an agarose gel, and DNA was precipitated using 1:1 of isopropanol at 20 °C for 30 min at 15,000g. The DNA pellet was dissolved in the buffer needed for its following application.

The sequence of the generated ASW14 fragment is:

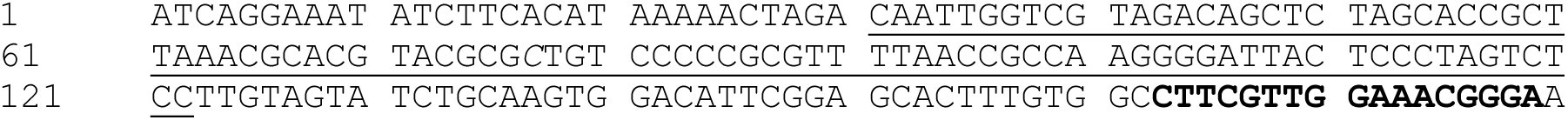
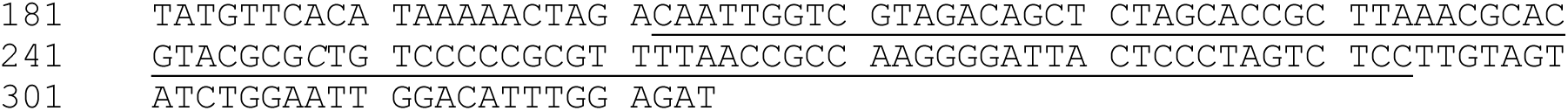

The Widom601 sequence underlined. Dyad base of the two CENP-A^Nuc^ indicated in italics. ASW14 with mutant B-box: The B-box of CTTCGTTGGAAACGGGA in bold of ASW14 was mutated to CAAAATTGGTAATTTTA in the B-box mutant α-satellite repeat dimer.

The 212 bp DNA fragment for the CCAN:CENP-A^Nuc^ complex (ASW6) has an additional 38 bp 5’ of an α-satellite repeat sequence (AS2) with a central 92 bp Widom601 sequence. ASW6 sequence is (B-box element in bold):

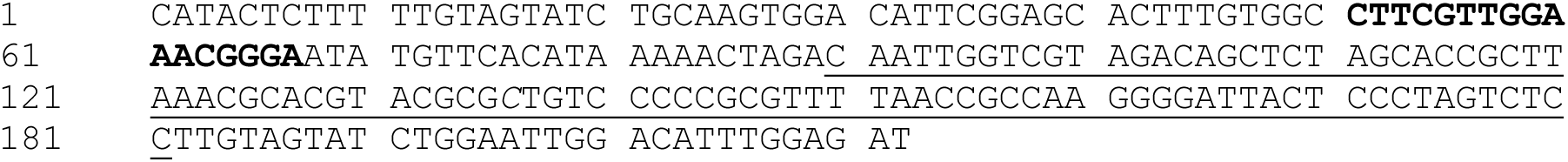

<u>AS2 (α-satellite repeat sequence from chromosome 2), also defined in Yatskevich et al.,</u> ^12^:

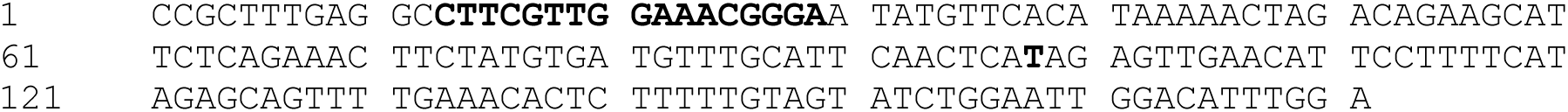

<u>α-satellite repeat sequence</u> as defined in Hasson et al., (Supplementary Figure 6a) ^13^.

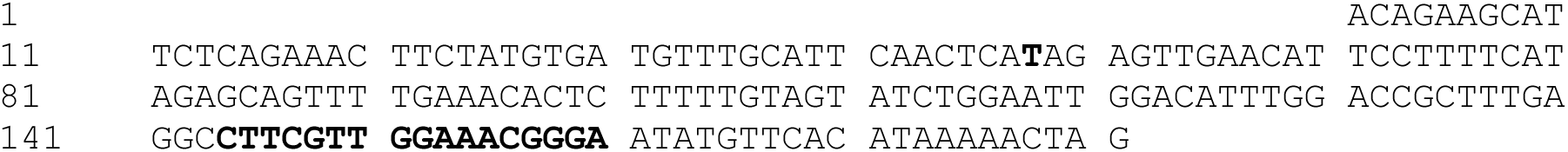

We define the start and end of the α-satellite repeat based on the positioning of the CENP-A nucleosome on the α-satellite repeat sequence from our 2.4 Å resolution cryo-EM structure ^12^. Specifically, we numbered the 3’ end of the repeat (nucleotide position #171) as the entry site of the CENP-A^Nuc^ DNA gyre. This then positions the CENP-A^Nuc^ dyad axis as nucleotide #98, and the 5’ entry into the DNA gyre as #25. Interestingly, the dyad axis of CENP-A^Nuc^ at nucleotide #98 maps to the centre of the 19 bp palindrome of human α-satellite repeats (TTCAACTCA**T**AGAGTTGAA), described in the 1978 papers first reporting primate α-satellite sequences ^54,55^. Our definition of the α-satellite repeat is 50 bp 5’ to that defined by Hasson et al., (2013) ^13^.

### CCAN:DNA complex assembly

CCAN:DNA complexes were assembled in CCAN buffer (300 mM NaCl, 20 mM Hepes pH 7.8, 0.5 mM TCEP) as follows. 6 µM CCAN was incubated at 22 °C for 30 mins before addition of an equal volume of 6 µM 171 bp (AS2 α-satellite repeat ^12^) DNA (AS2 described in ^12^), and incubated for a further 30 mins at at 22 °C, resulting in a final CCAN:DNA concentration of 3 µM and a final volume of 100 µL. The CCAN:DNA complex was then centrifuged at 10 °C in a table-top centrifuge for 10 min. To stabilize complexes for cryo-EM analysis, on-column crosslinking was performed.

### On-column crosslinking of CCAN:DNA

An Agilent 500 column coupled to a MicroAkta was pre-equilibrated in CCAN buffer. 100 µL 2.5 mM BS(PEG)_5_ (ThermoFisherScientific) was injected into the system for 1 mL at a flow rate of 0.07 µL/min. The CCAN:DNA complex was then injected into the column and eluted at the same flow rate into a 96-well plate in 50 μL fractions. Each plate well was supplemented with 5 µL 1 M Tris.HCl buffer pH 7.4 to quench the crosslinker during elution. Fractions containing CCAN:DNA were pooled, buffer exchanged into 150 mM NaCl, 20 mM Hepes pH 7.8, 0.5 mM TCEP and concentrated to 1.5 mg/mL (Amicon 100 kDa, 0.5 mL). Following concentration, CCAN:DNA complexes were again centrifuged at 10 °C in a table-top centrifuge for 10 min.

### CENP-A mono-nucleosome (ASW6) and CENP-A di-nucleosome (ASW14) reconstitution

The CENP-A histone octamer was prepared as described previously ^12^. Either ASW6 or ASW14 DNA were desalted into buffer B (20 mM HEPES pH 7.5, 2 M KCl). DNA at 8 μM was mixed with the CENP-A histone octamer at molar ratios of 1.1:1 protein:DNA for mono-CENP-A^Nuc^ and 2.1:1 protein:DNA for di-CENP-A^Nuc^. The histone octamers were then wrapped with DNA using the gradient dialysis method starting from 2 M KCl (in 20 mM HEPES pH 7.5, 1 mM EDTA) to 300 mM KCl over 16 h at 22 °C. The mixture was then dialysed against 20 mM HEPES pH 7.5, 300 mM KCl, 1 mM EDTA for a further 5 h. The wrapped mono-CENP-A and di-CENP-A nucleosomes were assessed by agarose gel and stored at 4 °C.

### Assembly of the CCAN:CENP-A^Nuc^ (ASW6 DNA) complex

CENP-A^Nuc^ and CCAN components (CENP-C^N^, CENP-LN, CENP-HIKM, CENP-TWSX, and CENP-OPQUR^ΔN^) were mixed at a 1:1 molar ratio at 5 μM in PBS buffer and TCEP. The sample was injected into an Agilent Bio SEC-5 1000A 4.6×300 column in CCAN gel filtration buffer (20 mM HEPES pH 7.8, 300 mM NaCl, 1 mM TCEP). The peak fractions were analysed using a 4-12% SDS-PAGE gel, stained with Instant Blue Coomassie Stain. To prepare the complex for structural analysis, on-column cross-linking using BS(PEG)_5_ (ThermoFisherScientific) was performed. 100 μl of 2.5 mM BS(PEG)_5_ was injected into an Agilent Bio SEC-5 1000A 4.6×300 column (equilibrated in PBS buffer) at a flow rate set of 0.07 µL/min, then 0.2 mL buffer, after which the CCAN:CENP-A^Nuc^ sample in (100 μl) was applied and eluted in PBS buffer. The sample was eluted into a 96-well plate in 50 μL fractions. Each plate well was supplemented with 5 µL 1 M Tris.HCl buffer pH 7.4 to quench the crosslinker during elution. The peak fractions of the CCAN:CENP-A^Nuc^ samples were pooled and concentrated to 1.5 mg/mL for cryo-EM grid preparation.

### Assembly of the CCAN:di-CENP-A^Nuc^ (ASW14 DNA) complex

For assembly of the CCAN:di-CENP-A^Nuc^ (ASW14 DNA) complex, we used the same procedure as for the CCAN:mono-CENP-A^Nuc^ (ASW6 DNA) complex except we used 20 mM HEPES pH 7.8, 300 mM NaCl, 1 mM TCEP buffer in the cross-linking final gel filtration step.

### Assembly of the CCAN^ΔCENP-C^:di-CENP-A^Nuc^ (ScFv) (ASW14 DNA) complex

#### 1. Cross-linking di-CENP-A^Nuc^ and ScFv (di-CENP-A^Nuc^:ScFv)

Di-CENP-A^Nuc^ and ScFv were combined at a 1:2.3 molar ratio and dialysed into a buffer of 20 mM HEPES pH 7.5, 50 mM NaCl. Cross-linking of the di-CENP-A^Nuc^:ScFv complex was performed using a 24 mL Superose 6 column by injecting 500 μL of 2.5 mM BS(PEG)_5_ at a flow rate of 0.25 mL.min^-1^ for 7 mL before loading the CENP-A^Nuc^:ScFv complex onto the column at the same flow rate. The cross-linking was assessed using SDS-PAGE gel. The cross-linked fractions were pooled and concentrated to 10 μM.

#### 2. Assembly of cross-linked di-CENP-A^Nuc^:ScFv with CCAN

All CCAN components (omitting CENP-C^N^) were without DTT. The components were at a 1:1:1:1:1 molar ratio (CENP-OPQUR:CENP-LN:CENP-HIKM:CENP-TWSX:di-CENP-A^Nuc^:ScFv) with CCAN components at a concentration of 5 μM. The resultant complex was cross-linked on an Agilent Bio SEC-5 1000A 4.6×300 column. Collected fractions were concentrated to 1.5-2 mg/mL for cryo-EM grid preparation.

### CCAN displaces CENP-B from di-CENP-A^Nuc^-B-box

#### Di-CENP-A^Nuc^-CENP-B binding test

Di-CENP-A^Nuc^ and CENP-B were mixed at a 1:2 ratio at 5 μM final concentration and injected into a Superose 6 Increase 10/300 column (Cytiva). Appropriate fractions were collected and concentrated.

#### Di-CENP-A^Nuc^-CENP-B-CCAN binding test

An equal molar ratio of CCAN^ΔCENP-C^ components (CENP-LN, CENP-HIKM, CENP-TWSX, and CENP-OPQUR^ΔN^) was added to the di-CENP-A^Nuc^:CENP-B complex at a final concentration of 5 μM of di-CENP-A^Nuc^ and 10 μM CENP-B and then loaded onto an Agilent Bio SEC-5 1000A 4.6×300 column. The eluted fractions were visualized using a 4-12% SDS-PAGE gel, stained with InstantBlue Coomassie Stain. In control experiments to assess the elution volume of CENP-B alone, and the di-CENP-A^Nuc^:CENP-B complex, the individual samples were run on the same Agilent Bio SEC-5 1000A 4.6×300 column as the di-CENP-A^Nuc^:CENP-B:CCAN sample (**Supplementary Fig. 6d-i**).

### Mass photometry

To assess the oligomeric state of CCAN:di-CENP-A^Nuc^ and CCAN^ΔCENP-C^:di-CENP-A^Nuc^, 18 µL of sample at approximately 100 nM was applied to wells in silicone gaskets on clean coverslips (Refeyn, UK) on a OneMP instrument (Refeyn). Movies were imaged for 60 s using the AcquireMP software (Refeyn) and the results were processed using the DiscoverMP software. The data were analysed by using automatic Gaussian fitting. The contrast intensities of landing events due to interference scattering were converted to masses using a calibration performed with a standard set of protein oligomers (P1, Refeyn) under similar buffer conditions. Data events were exported and plotted in Prism (GraphPad).

### Cryo-EM grid preparation and data collection

#### CCAN:DNA sample vitrification

Immediately prior to plunge-freezing, CCAN:DNA samples were supplemented with n-octyl-β-D-glucopyranoside (Glycon-biochem) to a final concentration of 0.05%. Quantifoil 300 mesh copper R1.2/1.3 grids (Quantifoil Micro Tools) were rendered hydrophilic by glow discharge in a Edwards S150B glow discharger for 1 min 15 s, setting 6, 30-35 mA, 1.2 kV, 0.2 mBar (0.15 Torr). Subsequently, 3 µL sample was applied to the grids, which were then mounted in a Vitrobot IV (ThermoFisherScientific), blotted with Whatman Filter paper 1 (0-0.5 s waiting time, 2 s blotting time, -7 blotting force), and plunge frozen in liquid ethane maintained at a temperature of -170 °C.

#### CCAN:DNA data acquisition

10,628 micrograph movies were collected on a Titan Krios (ThermoFisherScientific) electron microscope, equipped with a K3 direct electron detector (Gatan) and energy filter (Gatan) with a slit width of 20 eV, operating at 300 keV with a flux of 25 e⁻/pixel/s, at a nominal magnification of 105k, yielding a pixel size of 0.725 Å/pixel and a total dose of 40e/Å^2^. Automated data acquisition was performed using EPU (ThermoFisherScientific) with aberration-free image shift enabled, and a defocus range of -0.8 to -2.2 µm in 0.2 µm increments.

#### CCAN:CENP-A^Nuc^ complex sample vitrification and cryo-EM data acquisition

To prepare the CCAN:CENP-A^Nuc^ complex for cryo-EM, 3.5 µL of sample at a concentration of 2 mg/mL was applied to glow-discharged Quantifoil R2/1 300 mesh copper grids (Quantifoil Micro Tools). Grids were blotted for 2 s at 4°C, 100% humidity, and a blotting force of -10, then plunge-frozen in liquid ethane using a Vitrobot Mark IV (ThermoFisherScientific). Data were collected on a Titan Krios microscope (ThermoFisherScientific) operated at 300 kV, equipped with a K3 direct electron detector (Gatan) and an energy filter (Gatan) with an energy slit of 20 eV. Images were acquired at a nominal magnification of 81,000×, corresponding to a pixel size of 1.072 Å. Movies were recorded at a dose rate of 15 e⁻/pixel/s with an exposure time of 3 s over 40 frames. Automated data collection was performed using EPU (ThermoFisherScientific) with aberration-free image shift (AFIS) enabled. The defocus range was set from -1.0 to -2.0 µm in 0.2 µm increments. A total of 50,781 movie frames were collected.

#### CCAN:di-CENP-A^Nuc^ complex sample vitrification and cryo-EM data acquisition

To prepare the CCAN:di-CENP-A^Nuc^ complex for cryo-EM, 3.5 µL of sample at a concentration of 2 mg.mL^-1^ was applied to glow-discharged Quantifoil R1.2/1.3 300 mesh copper grids (Quantifoil Micro Tools). Grids were blotted for 2 s at 4°C, 100% humidity, and a blotting force of -7, then plunge-frozen in liquid ethane using a Vitrobot Mark IV (ThermoFisherScientific). Data were collected on a Titan Krios microscope (ThermoFisherScientific) operated at 300 kV, equipped with a K3 direct electron detector (Gatan) and an energy filter (Gatan) with an energy slit of 20 eV. Images were acquired at a nominal magnification of 81,000×, corresponding to a pixel size of 0.928 Å. Movies were recorded at a dose rate of 25 e⁻/pixel/s with an exposure time of 1.3 s over 82 frames. Automated data collection was performed using EPU (ThermoFisherScientific) with aberration-free image shift (AFIS) enabled. The defocus range was set from -1.0 to -2.0 µm in 0.2 µm increments. A total of 28,703 movie frames were collected.

### Cryo-EM data processing

#### CCAN:DNA complex

Micrograph movies were imported into RELION ^45^ and motion-corrected using RELION’s own implementation of MotionCor2 ^56^, followed by contrast transfer function estimation using the RELION wrapper for CTFFIND4 ^57^. Particle picking was performed using the crYOLO general model and particles were extracted in RELION with a box size of 150 pixels. Particles were subsequently exported to cryoSPARC and subjected to 2D classification. Particles exhibiting clear secondary structure features were used to generate an *ab initio* reconstruction. The initial particle set was then subjected to three rounds of heterogeneous refinement against one true volume (the CCAN:DNA *ab initio* reconstruction) and three noise decoy volumes arising from *ab initio* reconstruction of particles with no obvious proteo-typic features. Particles accumulating in the CCAN:DNA class were selected for each subsequent round of refinement until few or no particles reclassified into the noise decoy classes. The cleaned particle set was then subjected to homogeneous refinement, followed by 3D variability analysis. Particles exhibiting the most continuous DNA and CCAN subunit density were re-extracted without binning in RELION and subjected to particle polishing and CTF refinement. Finally, polished and CTF-corrected particles were re-imported into cryoSPARC where a final round of homogeneous refinement was performed, yielding a 3D reconstruction of CCAN:DNA to an overall resolution of 3.54 Å. The resulting map was filtered according to local resolution, as estimated in cryoSPARC, and then used for structural model building (in COOT) ^58^ and real-space refinement in PHENIX ^59^.

#### CCAN:CENP-A^Nuc^ complex

For the CCAN:CENP-A^Nuc^ complex, movie frames were motion-corrected, followed by estimation of Contrast transfer function (CTF) parameters using the patch-based method in cryoSPARC ^60^. Particle picking was performed with crYOLO ^61^ using the general model, and particles were extracted with a box size of 400 pixels. Reference-free 2D classification was then conducted to clean the particle dataset. Classes exhibiting clear secondary structure features were selected for *ab-initio* reconstruction and heterogeneous refinement in cryoSPARC. One class with 1,525,520 particles showing nucleosome features was selected for non-uniform refinement, yielding a reconstruction of the CENP-A^Nuc^:CENP-C^N^ at 2.4 Å resolution. Another dominant class containing 3,815,026 particles was import into RELION for further 3D classification. Three classes displaying CCAN feature were selected for particle subtraction using a mask covering 3’ CENP-A^Nuc^ region. After an additional round of 2D and 3D classification on the recentered particle dataset, 325,201 well-aligned nucleosome particles were reverted to their original, un-subtracted particles for further 3D classification. This yielded a subset of 39,036 particles corresponding to the CCAN:CENP-A^Nuc^ complex, which was refined to 4.5 Å resolution using Refine3D followed by post-processing.

#### CCAN:di-CENP-A^Nuc^ complex

For the CCAN:di-CENP-A^Nuc^ complex, all movie frames were motion-corrected, followed by estimation of Contrast transfer function (CTF) parameters using the patch-based method in cryoSPARC. Particle picking was performed with crYOLO using the general model, and the particles were extracted with a box size of 400 pixels. Reference-free 2D classification was then conducted to clean the particle dataset, and classes exhibiting clear secondary structure features were selected for *ab-initio* reconstruction and heterogeneous refinement in cryoSPARC. Three well-populated classes with clear structural features were further selected for non-uniform refinement, CTF refinement and local refinement, resulting three high-resolution maps: CENP-A^Nuc^:ScFv (1:1), CENP-A^Nuc^:ScFv (1:2) and CCAN:DNA, resolved at 2.7 Å, 2.7 Å and 2.9 Å, respectively.

To dissect the CENP-A^Nuc^ association with CCAN, the CCAN:DNA data was exported to RELION ^45^ for 3D classification, which revealed diffuse density at both the 3’ and 5’ end of the DNA. To recover CENP-A^Nuc^ at each side, two masks were generated and to subtract the density outside either the 3’ or 5’ CENP-A^Nuc^ regions. After re-centering the particle on the 3’ or 5’ CENP-A^Nuc^, nucleosome density was resolved on both sides, as confirmed by 2D classification and 3D classification showing dominant classes with clear nucleosome views. These dominant classes were reverted to their original un-subtracted particles for further 3D classification, yielding two subsets: 9,661 particles of CCAN:3’ CENP-A^Nuc^ and 5,396 particles of CCAN:5’ CENP-A^Nuc^. These were refined to 9.5 Å and 15.5 Å respectively. Subsequent multibody refinement and post-process modestly improved the map quality of both the CCAN:DNA and CENP-A^Nuc^ modules. Finally, the maps of CCAN with 3’ and 5’ CENP-A^Nuc^, were combined to generate a composite map of the CCAN di-CENP-A^Nuc^ complex.

### Cryo-EM model building and refinement

For the CCAN:DNA structure, the PDB entry 7R5S was used as the initial model. The model was fitted into the cryo-EM map of CCAN:DNA, and the DNA wrapping around CENP-TWSX was guided by a predicted CCAN:DNA model generated using AlphaFold3, followed by manual correction in COOT and real-space refinement in PHENIX. For the CCAN:CENP-A^Nuc^ structure, the newly refined CCAN:DNA complex model was docked into the corresponding map, followed by docking of a 3’ CENP-A^Nuc^ using PDB entry 7YWX. The models were then combined in COOT and real-space refined in PHENIX. For the CCAN:di-CENP-A^Nuc^ model, a composite map combining CCAN:3’ CENP-A^Nuc^ and CCAN:5’ CENP-A^Nuc^ was used. The CCAN:DNA model was first fitted into the map, followed by docking of CENP-A^Nuc^ models (PDB: 6E0P) to the 3’ and 5’ nucleosomes, respectively. All models were then combined in COOT, and the linker DNA region was manually built and refined. Additional manual corrections were also performed in COOT. Figures were generated using ChimeraX ^62^.

PDB and cryo-EM maps have been deposited with RCSB and EMDB, respectively. Accession numbers are listed in **Supplementary Table 1**. Source data are provided with this paper. Correspondence and requests for materials should be addressed to David Barford.

## Data availability

PDB coordinates generated in this study have been deposited with RCSB under accession codes 9TAW [https://www.rcsb.org/structure/9TAW], 9TAX [[https://www.rcsb.org/structure/9TAX] and 9TAY [https://www.rcsb.org/structure/9TAY]. Cryo-EM maps generated in this study have been deposited with EMDB under accession codes EMD-55755 []

EMD-55756 []

EMD-55757 []

EMD-55758 []

EMD-55759 []

Previously published protein coordinates used in study: 7R5S [https://www.rcsb.org/structure/7R5S], 7YWX [https://www.rcsb.org/structure/7YWX], 6E0P [https://www.rcsb.org/structure/6E0P] and 1AOI [https://www.rcsb.org/structure/1AOI]. Accession numbers are also listed in **Supplementary Table 1**. Source data are provided with this paper. Correspondence and requests for materials should be addressed to David Barford.

## Acknowledgements

We are grateful to the LMB EM Facility and the UK’s national Electron Bioimaging Centre (eBIC) (under proposal BI37660 funded by the Wellcome Trust, MRC and BBSRC) for help with the EM data collection; J. Grimmett, T. Darling and I. Clayson for scientific computing; K. Turton for help with insect cell expression. We thank Andrea Musacchio, Marion Pesenti and Ingrid Vetter for sharing unpublished experimental observations. For the purpose of open access, the authors have applied a CC BY public copyright license to any Author Accepted Manuscript version arising.

## Author Contributions

The project was conceived by D.B. Experiments were performed by C.Y, K.W.M., J.Y., Z.Z., D.B. The paper was written by D.B. with input from all other authors.

---

**Supplementary Fig. 1.**
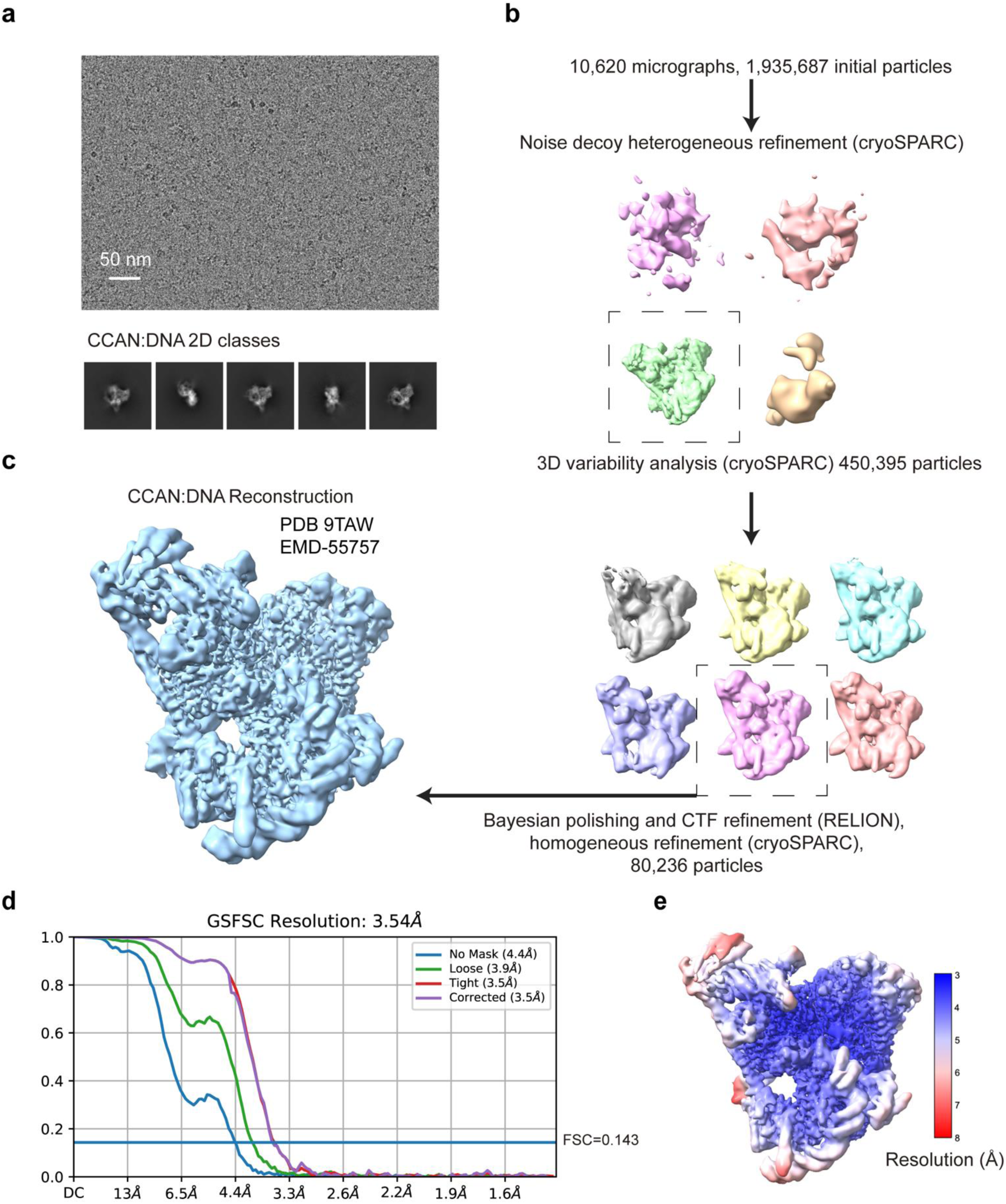
Cryo-EM data and processing workflow for the CCAN:DNA complex. **a**, Representative cryo-electron micrograph of 10,628 collected and 2D class averages for CCAN:DNA. **b**, Cryo-EM processing workflow. **c**, 3D reconstruction. A sharpened map is shown in Fig. 1c. **d**, FSC curves. **e**, Cryo-EM reconstruction colour-coded according to local resolution.

**Supplementary Fig. 2.**
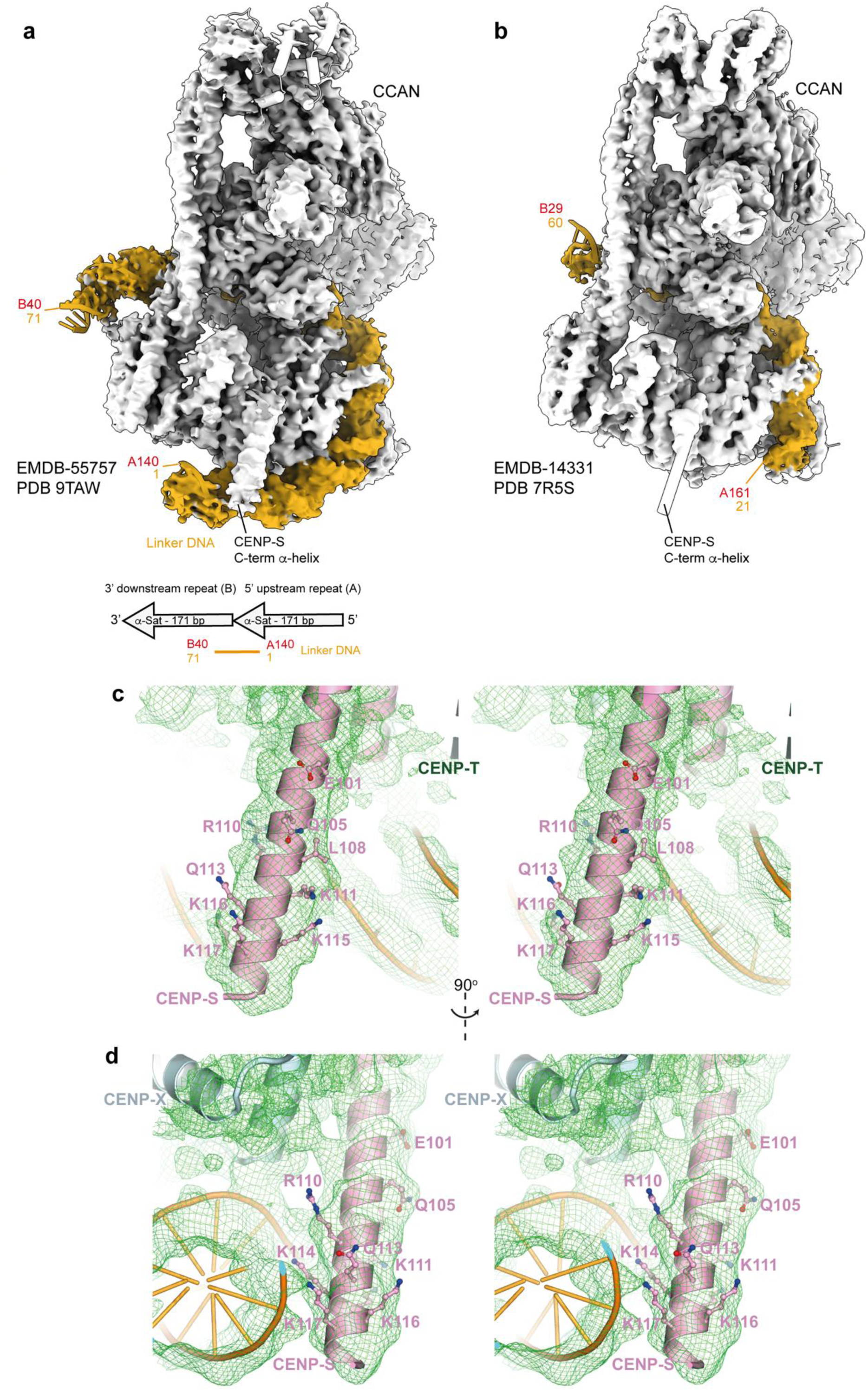
Cryo-EM density fits for DNA and CENP-S α-helix of the CCAN:DNA complex. **a**, Cryo-EM map of CCAN:DNA complex with CCAN density in grey and DNA density in orange. 70 bp of DNA are resolved in density. A140 and B40 refer to the positions on a dimeric α-satellite repeat sequence shown below and as defined in Methods and Fig. 3. **b**, Cryo-EM map of previous CCAN:DNA complex using 54 bp DNA which has 40 bp ordered ^12^. Base pair numbering matches (**a**). **c** and **d**, Orthogonal stereo views of cryo-EM density map for the ordered C-terminal α-helix of CENP-S. Residues shown in Fig. 2c are indicated. View in (**c**) as in Fig. 2c.

**Supplementary Fig. 3.**
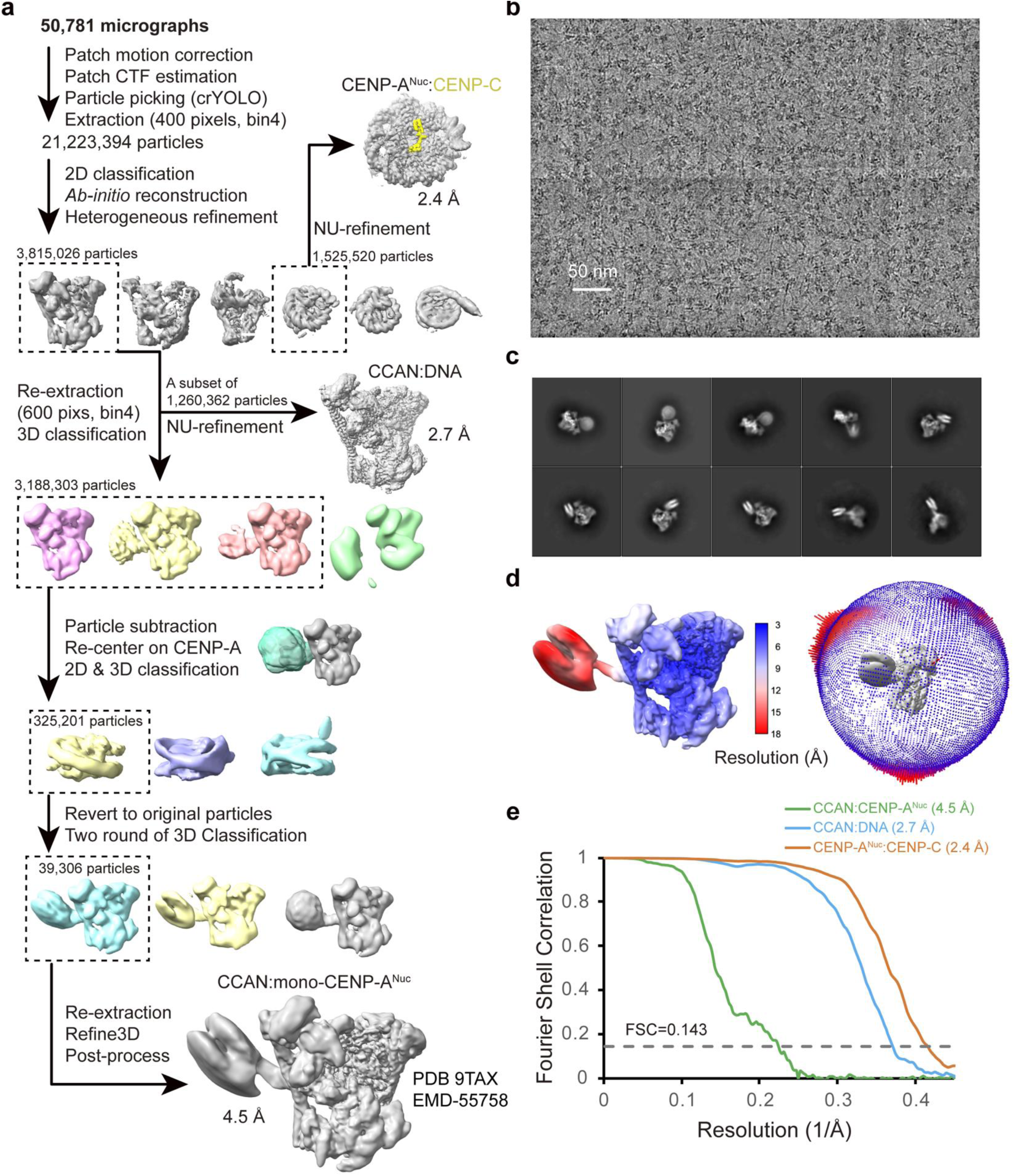
Cryo-EM images and processing workflow for CCAN:mono -CENP-A^Nuc^ complex. **a**, Cryo-EM processing workflow. **b**, Representative cryo-electron micrograph of 50,781 collected. **c**, 2D class average gallery. **d**, Cryo-EM reconstruction colour-coded according to local resolution. **e**, FSC curves. The CCAN:DNA coordinates and map were deposited with codes PDB ID 28OP and EMD-56683. The CENP-A^Nuc^:CENP-C coordinates and map were not deposited in PDB/RCSB with this study.

**Supplementary Fig. 4.**
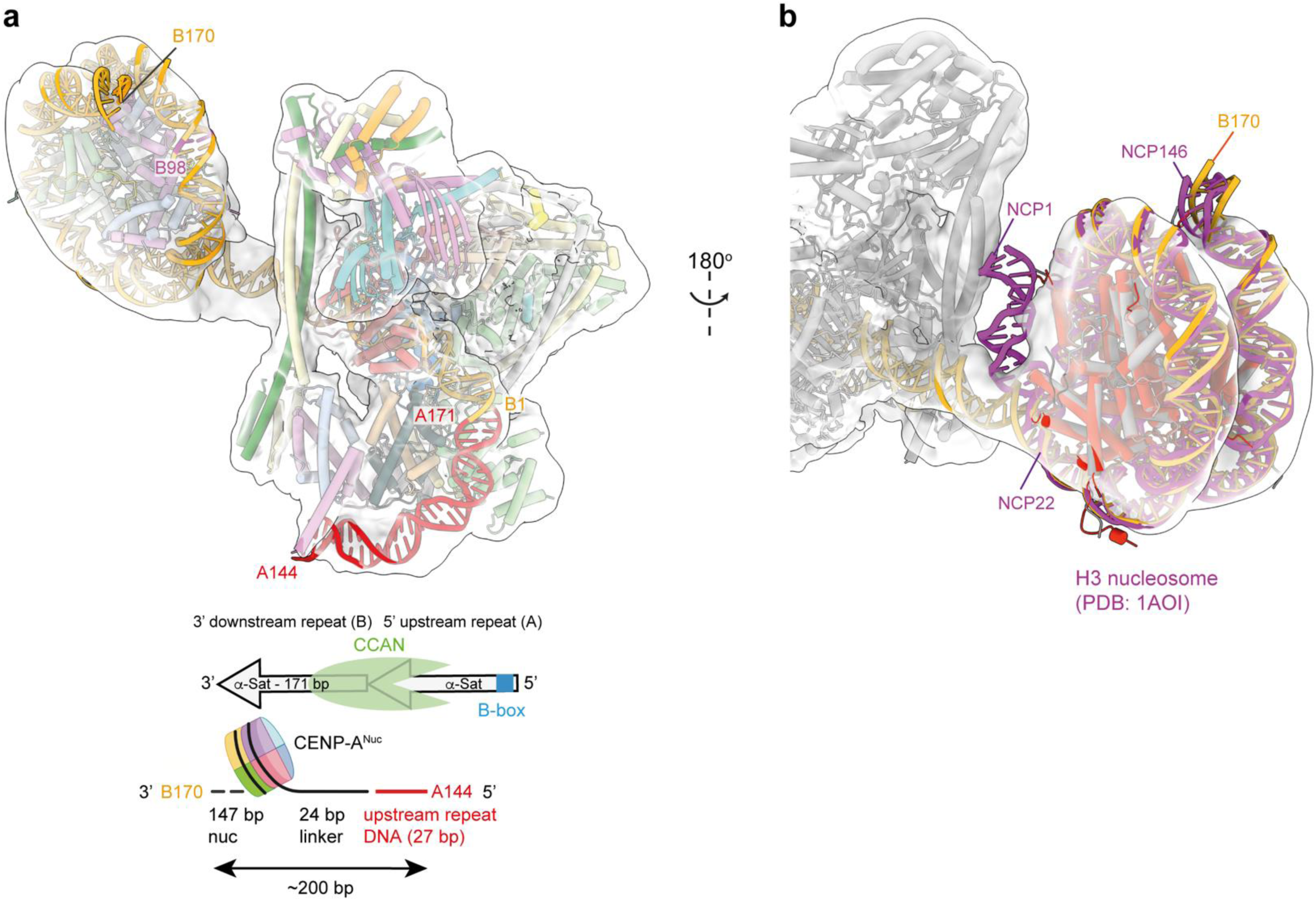
Cryo-EM density fits for DNA of the CCAN:CENP-A^Nu c^ complex. **a**, Cryo-EM map of CCAN:CENP-A^Nuc^ complex with CCAN density in grey and DNA density in red (5’ upstream repeat (A)) and orange (3’ downstream repeat (B)). A144 and B144 refer to the positions on a dimeric α-satellite repeat sequence shown in the schematic below and as defined in Methods. B98 indicates the dyad axis of CENP-ANuc. **b**, Superimposition of a canonical H3 nucleosome (NCP) (DNA in magenta, histones in red) onto the CENP -A nucleosome (DNA in orange, histones in grey). This illustrates that ∼20 bp of CENP-A^Nuc^ is unwrapped as the DNA gyre enters the CCAN DNA-binding tunnel.

**Supplementary Fig. 5.**
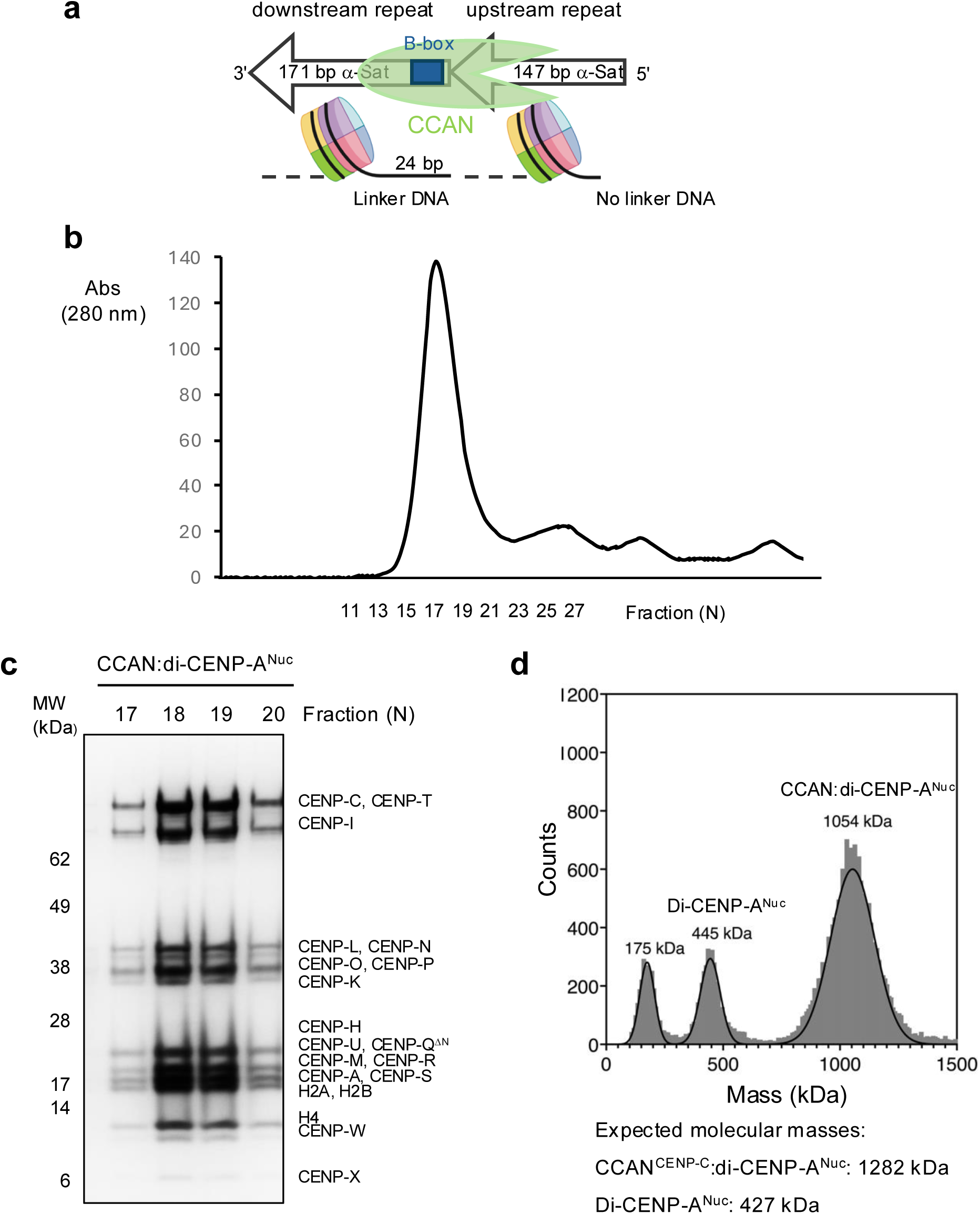
Reconstitution of CCAN:di-CENP-A^Nu c^ complex. **a**, Schematic of CCAN:di-CENP-A^Nuc^ complex showing α-satellite repeat dimer used in this study. The 3’ (downstream) repeat is 171 bp whereas the 5’ upstream repeat is 147 bp, i.e. without linker DNA. This design ensures only one CCAN-binding linker DNA per α-satellite repeat dimer. **b** and **c**, Size exclusion chromatogram (b) and associated SDS PAGE gel (c) of purified CCAN:di-CENP-A^Nuc^ complex. **d**, iSCAT data for CCAN:di-CENP-A^Nuc^ complex showing a mixture of di-CENP-A^Nuc^ alone (445 kDa) and the CCAN:di-CENP-A^Nuc^ complex (1,054 kDa).

**Supplementary Fig. 6.**
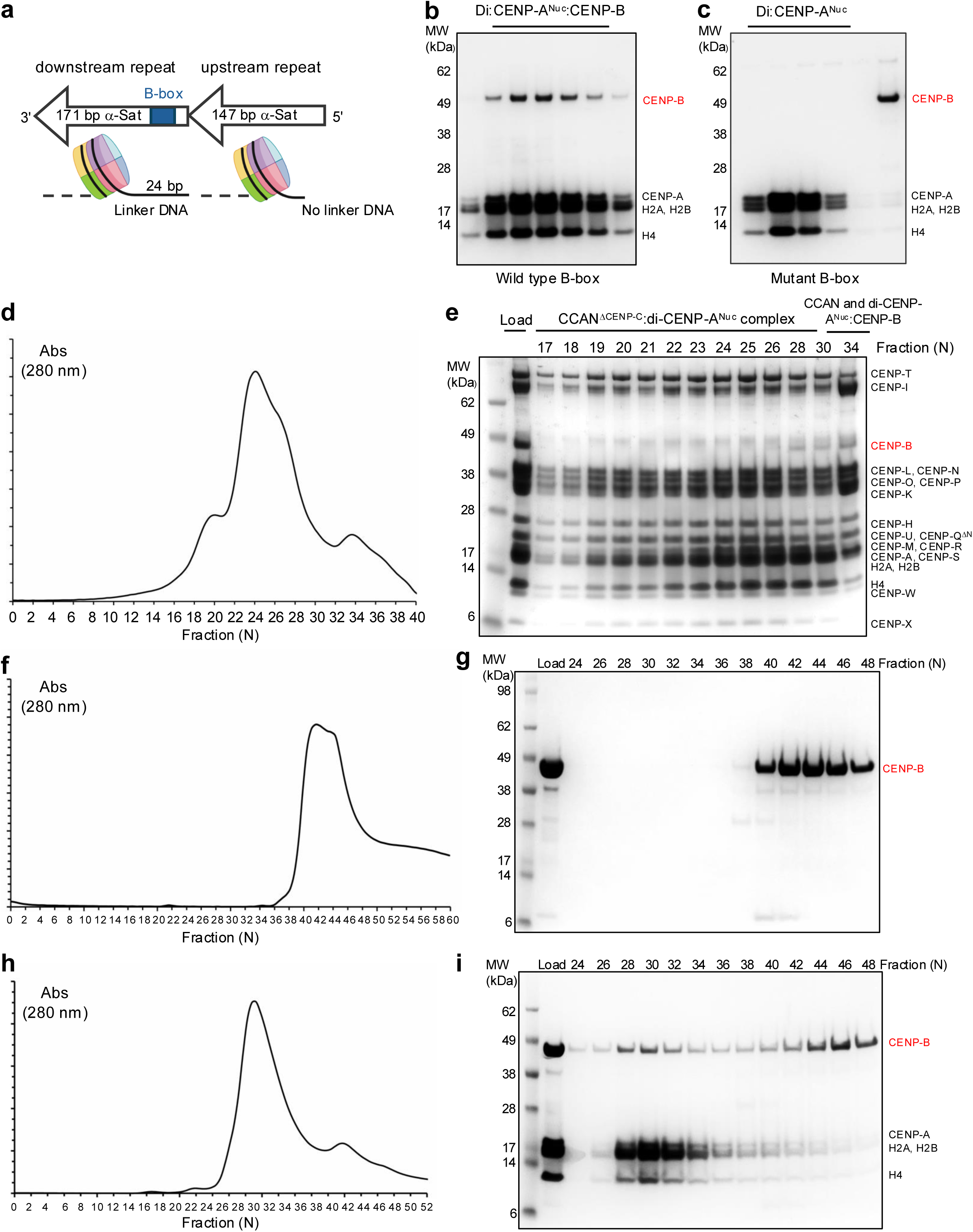
CCAN association with linker DNA displaces CENP -B bound to linker DNA B-box. **a**, Schematic of the CCAN:di-CENP-A^Nuc^ complex showing α-satellite repeat dimer used in this study with B-box in blue. SDS PAGE gels of: **b**, di-CENP-A^Nuc^:C ENP-B complex (with wild type B-box) and **c**, showing CENP-B does not bind to di-CENP-A^Nuc^ with a mutated B-box. **d**, and **e**, Size exclusion chromatogram (d) and associated SDS PAGE gel (e) showing that CCAN binding to di-CENP-A^Nuc^ displaces CENP-B in a load sample of di-CENP-A^Nuc^, CCAN and CENP-B (run on Agilent SEC-5 1000A 4.6×300 column). **f**, and **g**, Control SEC (f) and associated SDS PAGE gel (g) of CENP -B protein run on the same Agilent column. **h**, and **i**, Size exclusion chromatogram (h) and associated SDS PAGE gel ( i) of di-CENP-A^Nuc^:C ENP-B complex on the same Agilent column. CENP-B in lanes 28-34 in panel (e) is from di-CENP-A^Nuc^:C ENP-B complex (compare with panel (i)).

**Supplementary Fig. 7.**
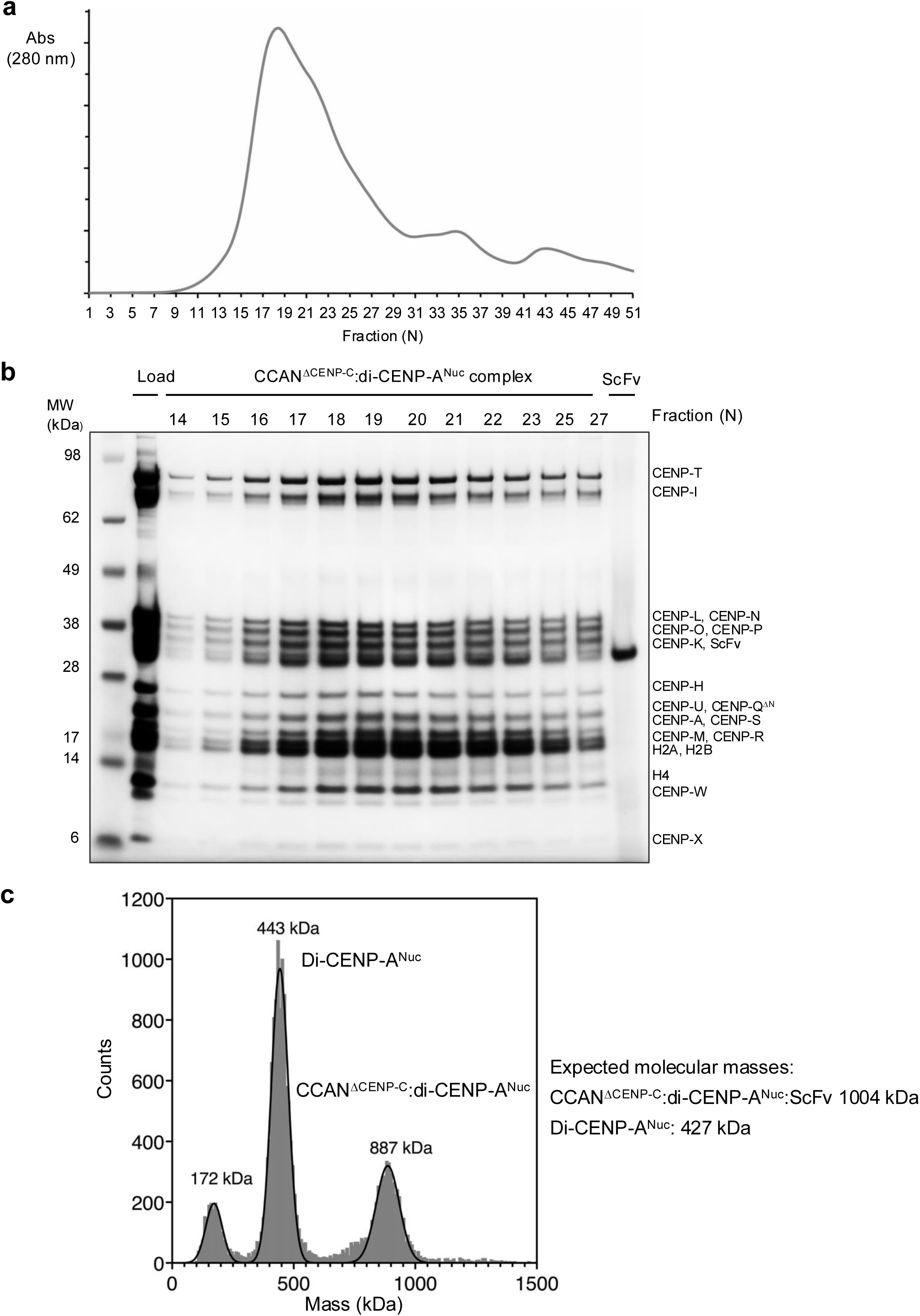
Reconstitution of CCAN^DCENP-C^:di-CENP-A^Nuc^ complex for cryo-EM. **a** and **b**, Size exclusion chromatogram (A) and associated SDS PAGE gel (B) of purified CCAN^ΔCENP-C^:di-CENP-A^Nuc^:ScFv complex. **c**, iSCAT data for CCAN^ΔCENP-C^:di-CENP-A^Nuc^:ScFv complex.

**Supplementary Fig. 8.**
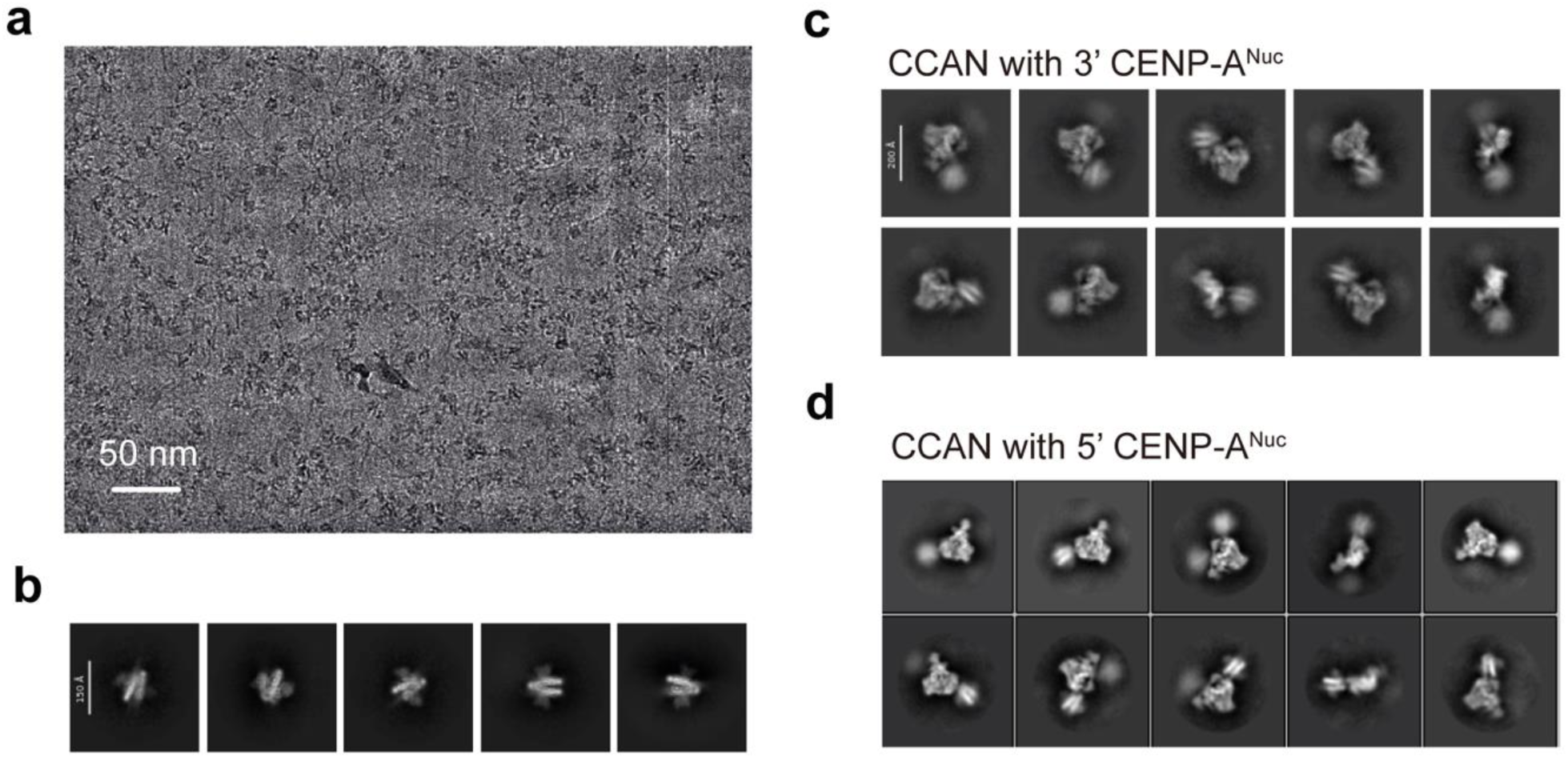
Cryo-EM data for the CCAN:di-CENP-A^Nuc^ complex. **a**, Representative cryo-electron micrograph of 28,703 collected. **b**, 2D class averages of CENP-A^Nuc^. **c**, 2D class averages of CCAN with 3’ CENP-A^Nuc^. **d**, 2D class averages of CCAN with 5’ CENP-A^Nuc^.

**Supplementary Fig. 9.**
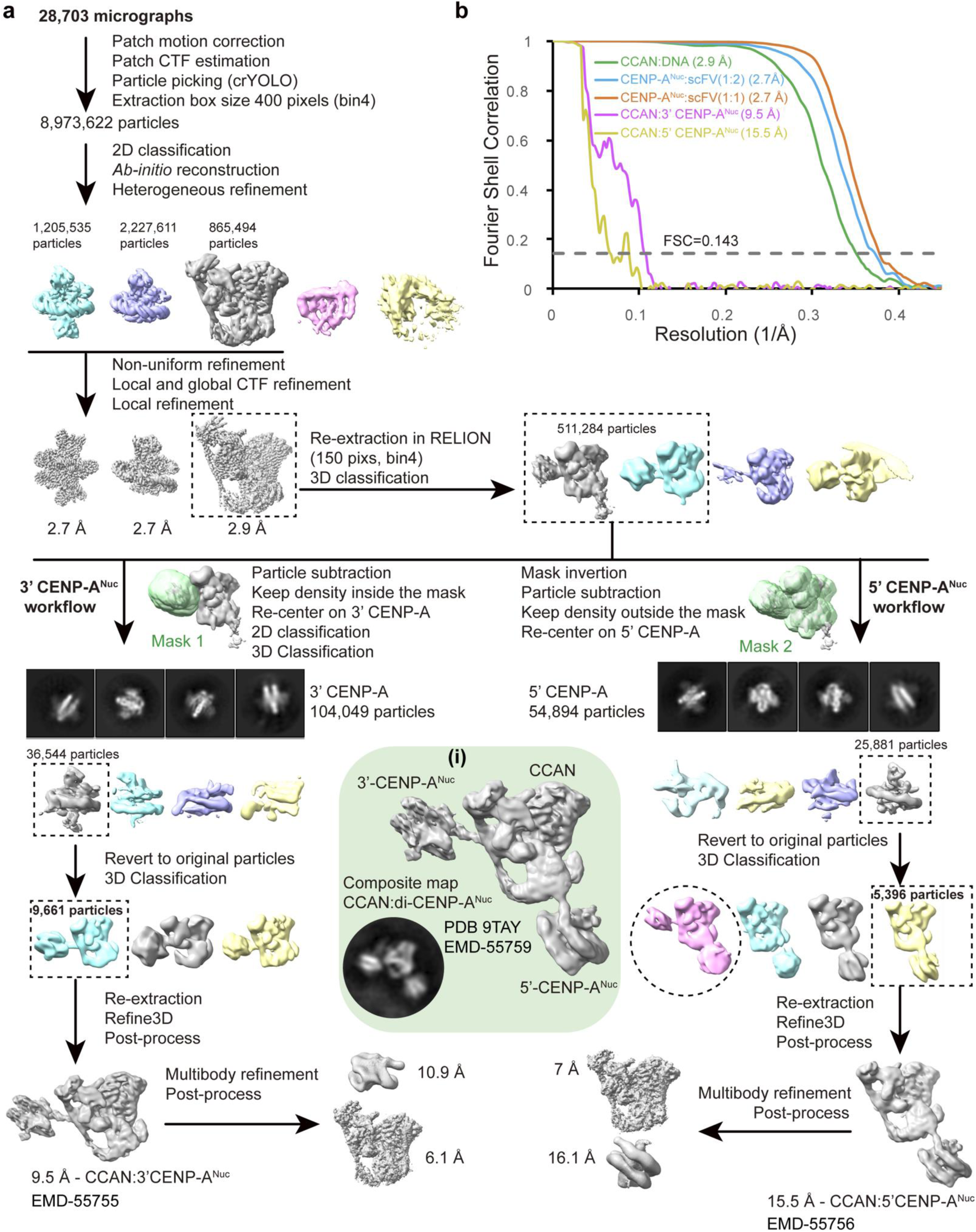
Cryo-EM workflow for CCAN:di-CENP-A^Nu c^ complex reconstruction. **a**, Cryo-EM processing workflow with insert **(i)** showing the composite map and 2D class average. **b**, FSC curves. Cryo-EM maps and coordinates for the CCAN:DNA, CENP-A^Nuc^:ScFv (1:2) and CENP-A^Nuc^:ScFv (1:1) complexes were not deposited with this study.

**Supplementary Fig. 10.**
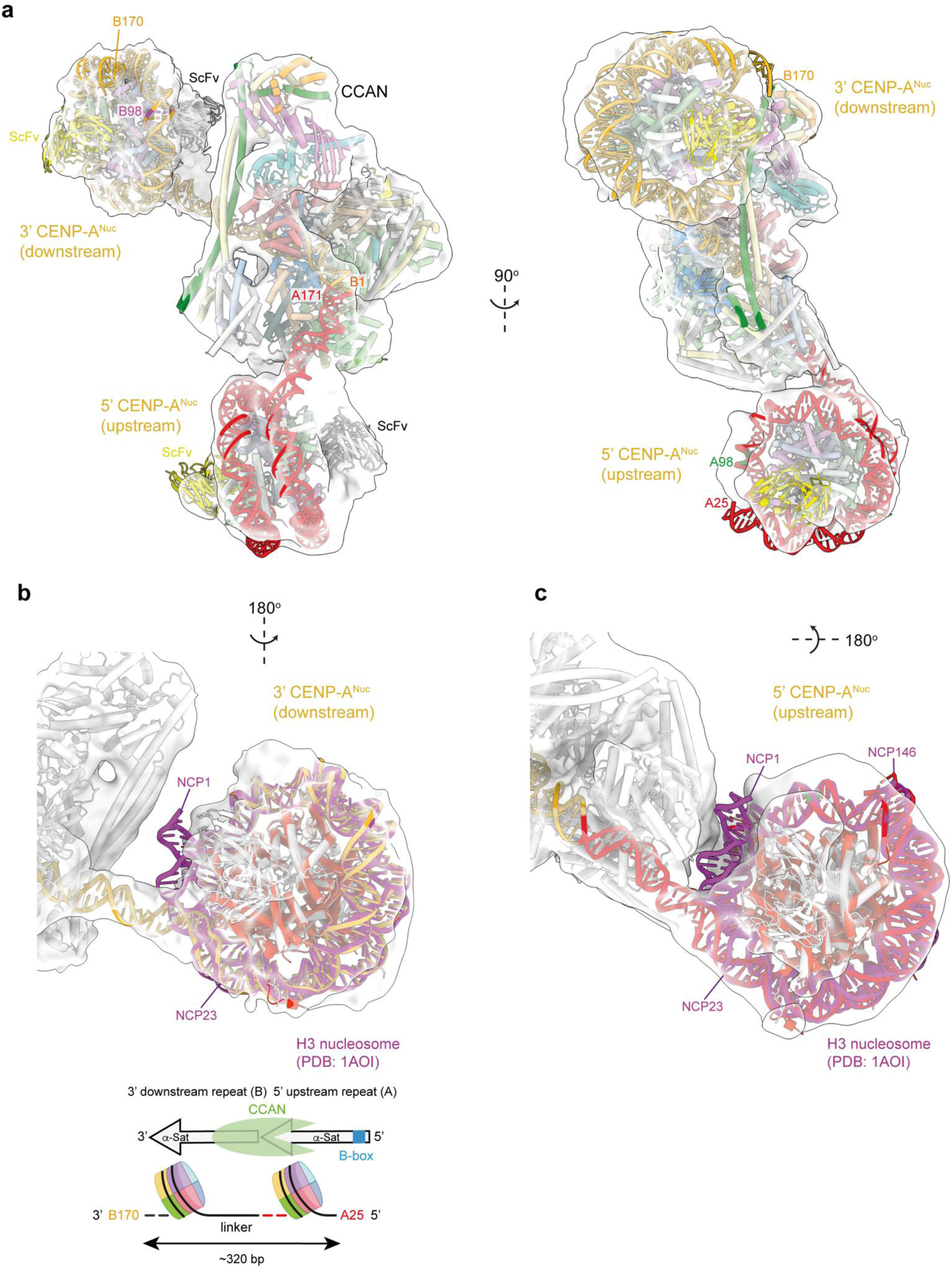
Cryo-EM density fits for DNA of the CCAN:di-CENP-A^Nuc^ complex. **a**, Cryo-EM map of CCAN:di-CENP-A^Nuc^ complex with CCAN density in grey and DNA density in red (5’ upstream repeat (A)) and orange (3’ downstream repeat (B)). A144 and B144 refer to the positions on a dimeric α-satellite repeat sequence shown in the schematic below and as defined in Methods. B98 indicates the dyad axis of CENP-ANuc. **b**, Superimposition of a canonical H3 nucleosome (NCP) (DNA in magenta, histones in red) onto the CENP-A nucleosome (DNA in orange, histones in grey). This illustrates that ∼20 bp of CENP-A^Nuc^ is unwrapped as the DNA gyre enters the CCAN DNA-binding tunnel.

**Extended Data Table 1.**
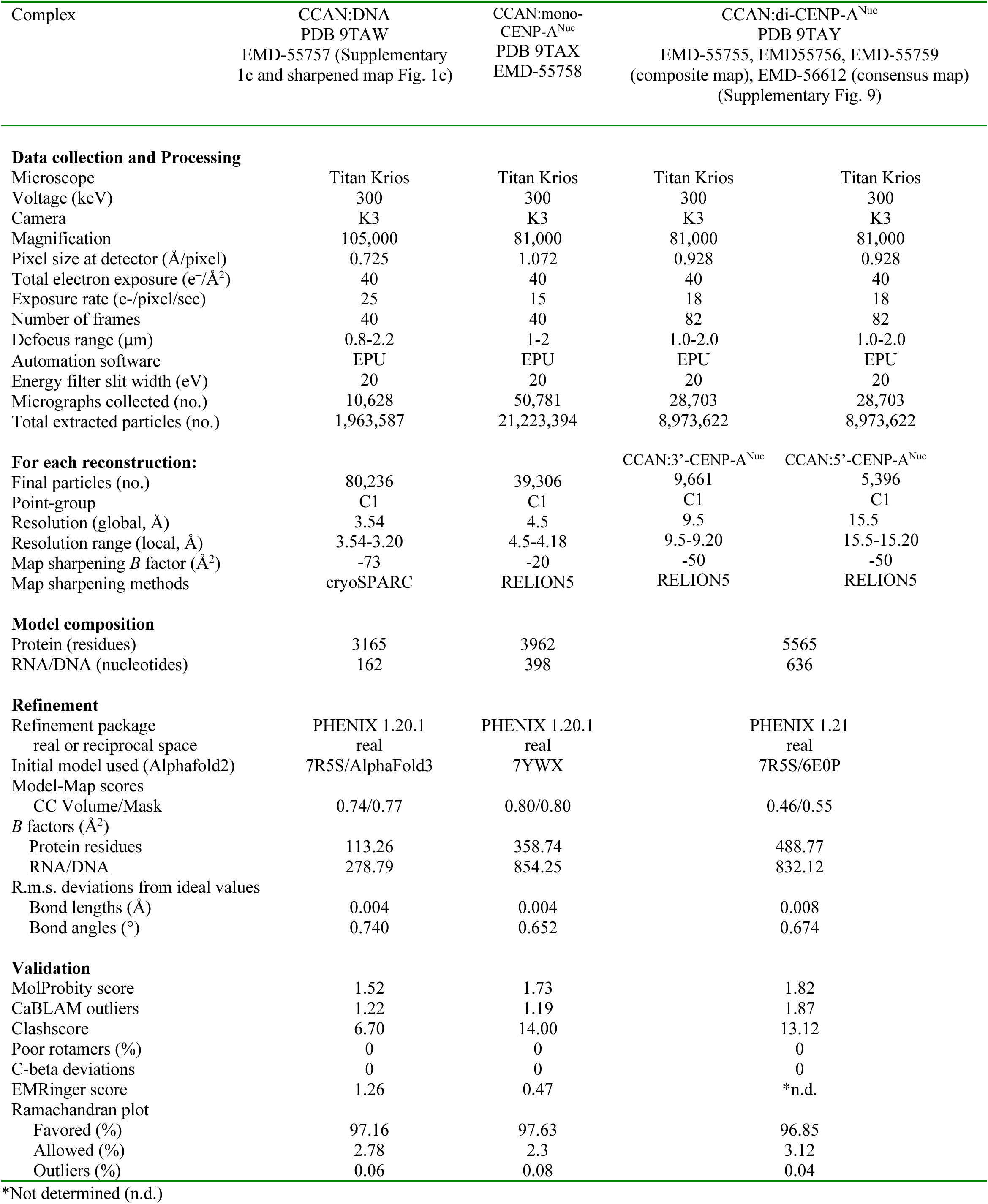
Cryo-EM data collection, refinement, and validation statistics.

